# Protein-Metabolite Interactomics Reveals Novel Regulation of Carbohydrate Metabolism

**DOI:** 10.1101/2021.08.28.458030

**Authors:** Kevin G. Hicks, Ahmad A. Cluntun, Heidi L. Schubert, Sean R. Hackett, Jordan A. Berg, Paul G. Leonard, Mariana A. Ajalla Aleixo, Aubrie Blevins, Paige Barta, Samantha Tilley, Sarah Fogarty, Jacob M. Winter, Hee-Chul Ahn, Karen N. Allen, Samuel Block, Iara A. Cardoso, Jianping Ding, Ingrid Dreveny, Clarke Gasper, Quinn Ho, Atsushi Matsuura, Michael J. Palladino, Sabin Prajapati, PengKai Sun, Kai Tittmann, Dean R. Tolan, Judith Unterlass, Andrew P. VanDemark, Matthew G. Vander Heiden, Bradley A. Webb, Cai-Hong Yun, PengKai Zhap, Christopher P. Hill, Maria Cristina Nonato, Florian L. Muller, Daniel E. Gottschling, James E. Cox, Jared Rutter

## Abstract

Metabolism is highly interconnected and also has profound effects on other cellular processes. However, the interactions between metabolites and proteins that mediate this connectivity are frequently low affinity and difficult to discover, hampering our understanding of this important area of cellular biochemistry. Therefore, we developed the MIDAS platform, which can identify protein-metabolite interactions with great sensitivity. We analyzed 33 enzymes from central carbon metabolism and identified 830 protein-metabolite interactions that were mostly novel, but also included known regulators, substrates, products and their analogs. We validated previously unknown interactions, including two atomic-resolution structures of novel protein-metabolite complexes. We also found that both ATP and long-chain fatty acyl-CoAs inhibit lactate dehydrogenase A (LDHA), but not LDHB, at physiological concentrations *in vitro*. Treating cells with long-chain fatty acids caused a loss of pyruvate/lactate interconversion, but only in cells reliant on LDHA. We propose that these regulatory mechanisms are part of the metabolic connectivity that enables survival in an ever-changing nutrient environment, and that MIDAS enables a broader and deeper understanding of that network.

## Main Text

Metabolites are the small molecule substrates, intermediates, and end products of metabolic pathways, and their physical interactions with proteins are among the most common and important interactions in biology (Fig. 1A). Metabolites are not only the chemical ingredients of metabolic reactions, but also enact enzyme regulation within and between metabolic pathways. Such regulatory interactions enable the maintenance of metabolic homeostasis in an environment where the availability and identity of nutrients are constantly changing. Despite their central importance, progress towards comprehensive identification of protein-metabolite interactions (PMIs) has been limited and sporadic. Unlike interactions involving other biological molecules, where we have robust and generalizable approaches such as coimmunoprecipitation for identifying protein-protein interactions (*1*) and chromatin immunoprecipitation for elucidating protein-DNA interactions (*2, 3*), widely applicable strategies to detect PMIs are lacking. Although some progress has been made recently (*4*), the very nature of many of these biologically important interactions presents a major hurdle to their identification. For example, to maximize the regulatory potential of protein binding, metabolites frequently interact with proteins with an affinity that is sufficiently weak to approximate their dynamic cellular concentrations—often high micromolar to low millimolar. Therefore, we developed the highly sensitive MIDAS (<u>M</u>ass spectrometry <u>I</u>ntegrated with equilibrium <u>D</u>ialysis for the discovery of <u>A</u>llostery <u>S</u>ystematically) platform to enable the systematic discovery of PMIs, including those low-affinity interactions that mediate cellular metabolic homeostasis.

**Figure 1.**
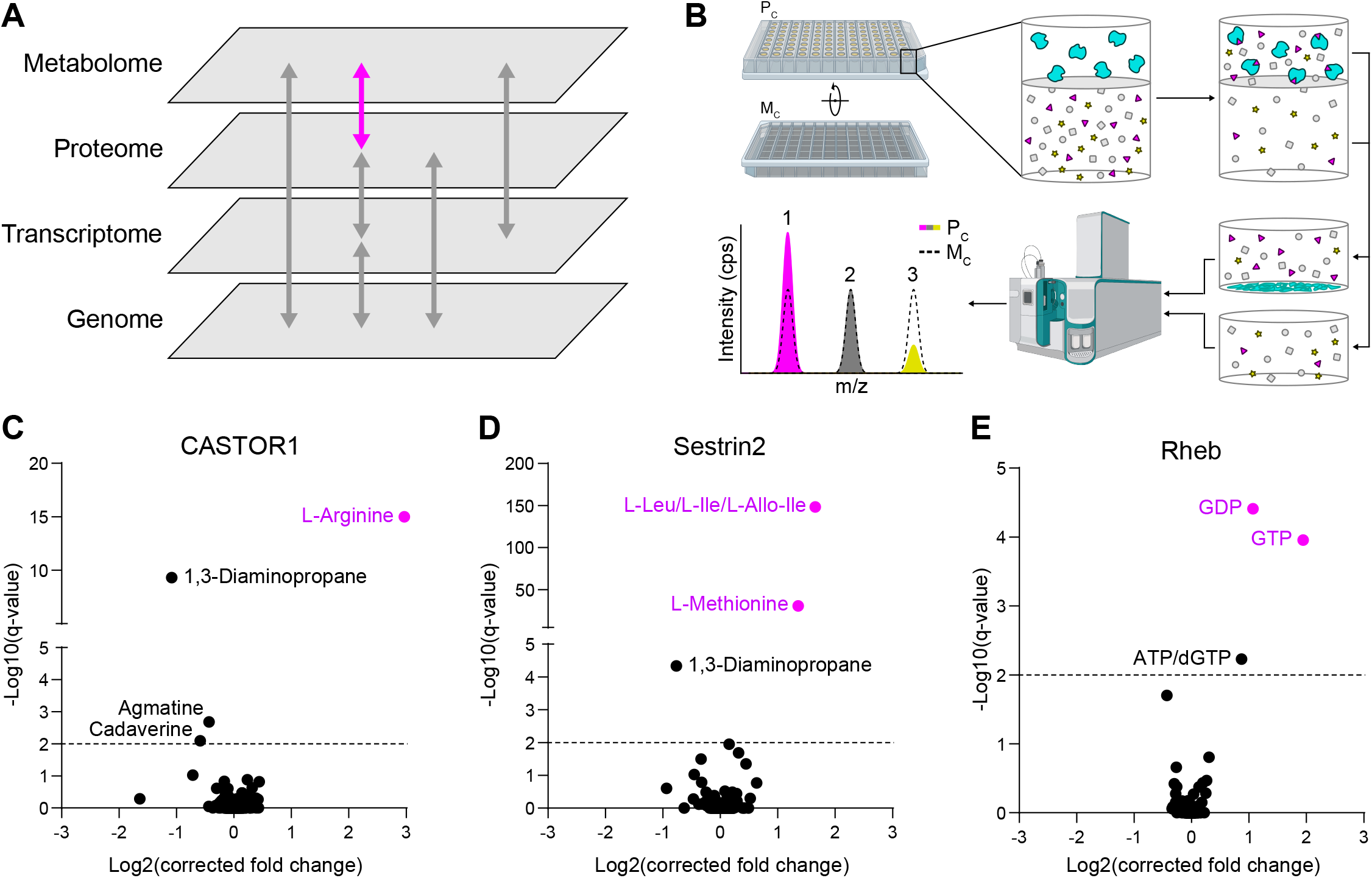
MIDAS is a platform for the systematic discovery of protein-metabolite interactions. **(A)** Biological systems are organized into domains of information (panes). Movement and interaction of information in and through these domains underlies biological function (arrows). The MIDAS platform provides protein-metabolite interactome (purple arrow) discovery. **(B)** The MIDAS platform is an equilibrium dialysis tandem mass spectrometry approach. (Top) Purified proteins (cyan) are loaded into the protein chamber (P_c_) and defined pools of metabolites into the metabolite chamber (M_c_), separated by a protein-impermeable membrane. The system is incubated to relative equilibrium. (Bottom) Proteins are removed, the P_c_ and M_c_ are sampled, and the relative abundance of metabolites from each chamber are quantified using FIA-MS. Interactions between proteins and metabolites are observed as an increase (1, magenta) or decrease (3, yellow) in integrated signal intensity in the P_c_ relative to the M_c_ (dotted peak). Metabolites that have equal integrated signal intensity in the P_c_ relative to the M_c_ (2, grey) are defined as non-interacting. **(C, D, E)** The mTORC1 regulators CASTOR, Sestrin2, and Rheb were screened using MIDAS. Known metabolite regulators are highlighted (magenta). Significant protein-metabolite interactions have a q-value < 0.01 (dotted line).

The MIDAS platform is built on the biophysical principle of equilibrium dialysis (*5*) (Fig. 1B). A purified protein is separated from a defined library of metabolites by a semi-permeable dialysis membrane that allows passage of the metabolites but not the protein. After incubation, the system achieves relative equilibrium, such that the concentration of free (i.e. non-interacting) metabolites becomes similar on both sides of the membrane. However, the total concentration of those metabolites that interact with the protein is specifically elevated in the protein-containing chamber (Fig. 1B-purple triangles). The protein is then removed by precipitation and the relative metabolite abundance is quantified by high-throughput flow injection analysis mass spectrometry (FIA-MS). Interestingly, there are also metabolites that are selectively depleted from the protein-containing chamber (Fig. 1B-yellow stars), which likely results from enzymatic conversion or from very high affinity interactions including covalent protein modifications.

The MIDAS metabolite library comprises 401 compounds that together represent a sizable fraction of the water-soluble, stable, FIA-MS-detectable, and commercially available human metabolome (Fig. S1A and Data S1). Due to intrinsic differences in chemical structure and ionization properties, not all metabolites could be analyzed using the same FIA-MS parameters. We profiled each metabolite individually for its optimal FIA-MS polarity and mobile phase pH ionization and detection conditions (Data S2), and, guided by these criteria, we divided the library into four pools for multiplexed analysis (Fig. S1B and Data S1). Next, we developed FIA-MS methods, optimized for each pool, that enabled rapid and robust quantitation of the constituent metabolites. Splitting metabolites into pools also allowed us to separate some isomers so that they could be independently quantified, solving a technical shortcoming of FIA-MS.

Having established the MIDAS methodology, we performed a pilot validation study using proteins with well-established metabolite interactions. We analyzed three human proteins that converge to regulate the mTORC1 kinase, an important nexus in growth factor signaling: CASTOR1, which binds arginine (*6*); Sestrin2, which binds leucine, isoleucine, and methionine with decreasing affinity (*7*); and Rheb, which is a GTPase that hydrolyses GTP to GDP (*8*) (Fig. S1C). In each case, the known ligands were the most enriched and statistically significant interactors detected: CASTOR1 enriched arginine (Fig. 1C); Sestrin2 enriched the leucine/isoleucine/allo-isoleucine isomer group and methionine (Fig. 1D); and Rheb enriched both its substrate, GTP, and its product, GDP, as well as the structually similar isomer group of ATP and deoxyGTP (Fig. 1E). Thus, MIDAS effectively identified known PMIs, including regulators as well as enzyme substrates and products.

The enzymes of glycolysis and related metabolic pathways are of particular interest for MIDAS analysis due to their importance in almost all human cells and the extent of known metabolite interactions. Therefore, we next used MIDAS to profile 33 human enzymes of central carbon metabolism, including enzymes of glycolysis, gluconeogensis, the tricarboxylic acid (TCA) cycle, and the serine biosynthetic pathway that emerges from glycolysis (Fig. S1C). In total, we identified 830 putative PMIs, the vast majority of which were previously unknown (Table S3). Unsupervised hierarchical clustering (Fig. 2A) and multidimensional scaling (Fig. 2E) of the entire PMI dataset demonstrated that structurally and functionally related proteins frequently have very similar metabolite interactions. For example, phosphoglycerate mutase (PGAM1/2), enolase (ENO1/2), fructose bisphosphatase (FBP1/2), and lactate dehydrogenase (LDHA/B) isoforms all clustered closely together. However, this was not observed across all enzyme isoforms nor would it be expected given the known role of enzyme isoforms to enable distinct biological regulation of pathways in different contexts. The PK-M1 isoform of pyruvate kinase was noticeably different from the PK-LR and PK-M2 isoforms, and the IDH2 and IDH3 isoforms of isocitrate dehydrogenase, which catalyze similar chemistry but are evolutionarily and structurally unrelated (*9*), exhibited distinct metabolite interactomes. Additionally, we observed clustering of multiple NAD(H)-dependent dehydrogenases: glyceraldehyde-3-phosphate dehydrogenase (GAPDH), LDHA, LDHB, mitochondrial malate dehydrogenase (MDH2), and 3-phosphoglycerate dehydrogenase (PHGDH). An analogous clustering of structurally and functionally related metabolites was also apparent, including nicotinamide-containing metabolites and flavin-adenine dinucleotide (Fig. 2B), phosphate-containing organic acids (Fig. 2C), and several nucleotide monophosphates (Fig. 2D).

**Figure 2.**
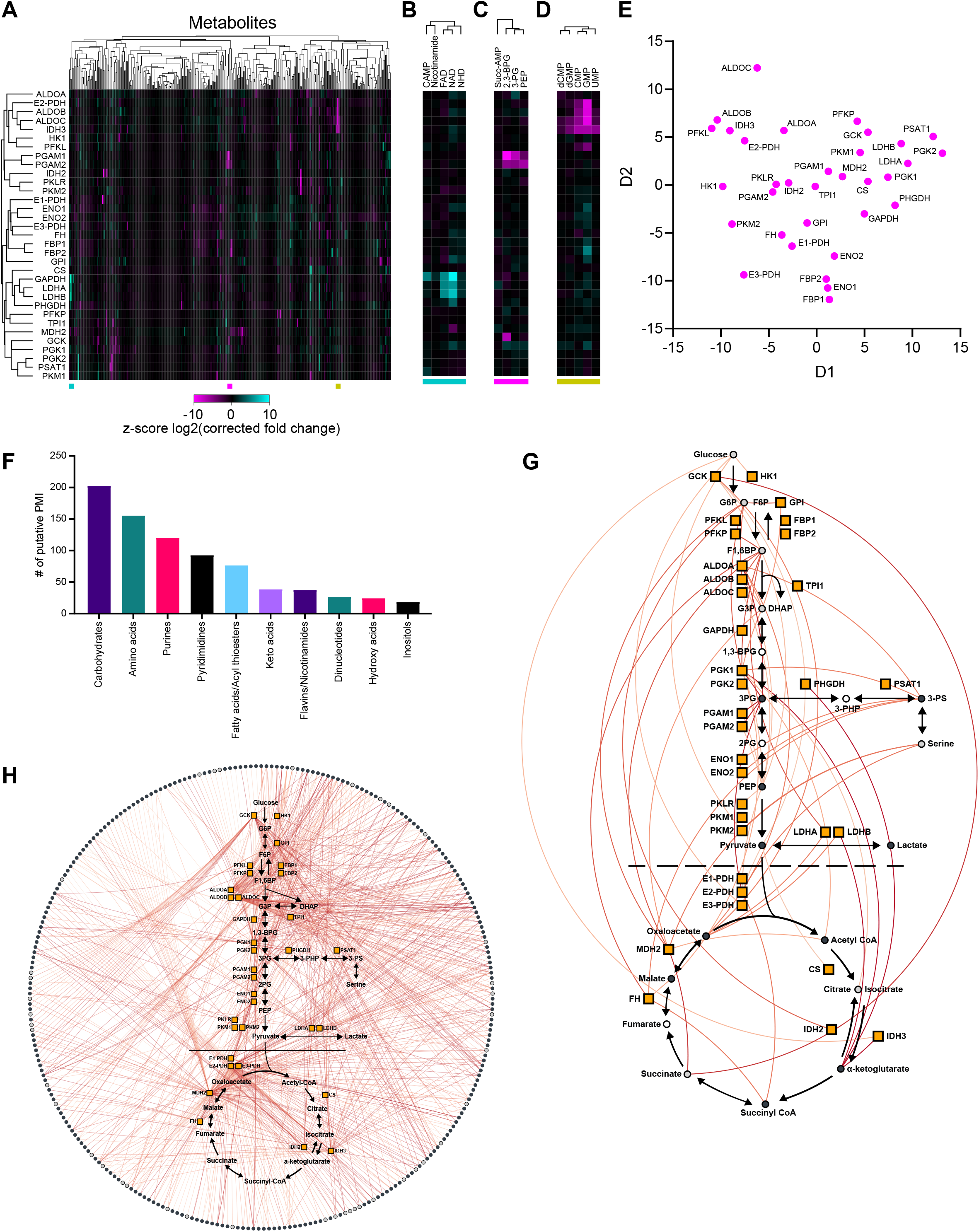
The protein-metabolite interactome of human central carbon metabolism. **(A)** Heatmap representation of the MIDAS protein-metabolite interactomes of 33 enzymes in central carbon metabolism. Heatmap values are the z-score log2(corrected fold change) for all metabolites in the MIDAS metabolite library on a per protein basis. Clustering was performed by one minus Pearson correlation. Positive (cyan) and negative (magenta) metabolite z-score log2(corrected fold change) have a maximum and minimum cut-off of 10 and –10, respectively. **(B, C, D)** Excerpt examples of metabolite clustering from (A). Colored bars (bottom) indicate the location of the extracted heatmaps from (A) (bottom). **(E)** Multidimensional scaling (MDS) of 33 enzymes in central carbon metabolism based on their MIDAS protein-metabolite interactomes. MDS distance values where generated from the z-score log2(corrected fold change) for all metabolites in the MIDAS metabolite library on a per protein basis. **(F)** The top ten metabolite sub-classes by total protein-metabolite interaction (PMI) count across 33 enzymes in human central carbon metabolism. Metabolite sub-classes were modified from HMDB chemical taxonomy sub-class. **(G and H)** Significant intra-pathway (G) and inter-pathway (H) interactions (colored lines) between metabolites (black circles) and 33 enzymes in central carbon metabolism (orange boxes) detected by MIDAS (plots generated in *Electrum*). Unique metabolites (dark grey circles), metabolite isoforms (light grey circles), metabolites not present in the library (open circles). Significant protein-metabolite interactions have a q-value < 0.01 and are colored by increasing significance, light orange to red.

Analysis of the 830 putative PMIs identified by the MIDAS platform showed that carbohydrates were the predominant class of protein-interacting metabolite across central carbon metabolism (Fig. 2F). This likely reflects both substrate/product relationships as well as the allosteric regulation of these enzymes, which are largely involved in carbohydrate metabolism, by upstream or downstream metabolites (i.e., feedforward and feedback regulation). The majority of non-carbohydrate PMIs included amino acids, nucleotides, and fatty acid derivatives. Such PMIs not only represent substrates and products of enzymes in these pathways, but suggest both intra- and inter-pathway regulation of central carbon metabolism (Fig. 2G). Notably, we also observed extensive interactions with metabolites outside of these pathways (Fig. 2H). Together, these data likely illustrate the integration of local and distal metabolic information on central carbon metabolism, which provides intermediates for most biosynthetic pathways in the cell.

We next selected a subset of PMI datasets from individual proteins in central carbon metabolism (see Fig. 2G) for deeper bioinformatic, biochemical and structural analyses. Enolase catalyzes the penultimate step in glycolysis, and the most enriched and most statistically significant interactor for both isoforms (ENO1 and ENO2) was phosphoserine (pSer, Fig. 3A). Intriguingly, pSer is an intermediate in the serine biosynthetic pathway, which diverges from glycolysis just upstream of enolase. pSer is subsequently converted into serine which in turn allosterically activates the M2 isoform of pyruvate kinase (*10*), the enzyme immediately downstream of enolase in the glycolytic pathway. Differential scanning fluorimetry (DSF), which measures the thermal stability of a target protein, showed that pSer (but not serine, phosphotyrosine, or phosphate) robustly stabilized both ENO1 (Kd_app_ = 1.38 mM) and ENO2 (Kd_app_ = 1.15 mM) (Fig. 3B), similar to their substrate 2PG (Kd_app_ = 0.298 mM and 0.289 mM, respectively). X-ray crystallography of the pSer-ENO2 complex found that pSer asymmetrically bound to the ENO2 dimer at one of the two active sites and partially overlapped with the 2PG phosphate binding site (Fig. 3C, D). Furthermore, pSer promoted an “open” active site conformation relative to the substrate bound complex observed as repositioning of loops 4 and 11 and alpha helices 7 and 11 (Fig. 3D). Surprisingly, pSer only weakly inhibited enolase activity as measured in vitro (Fig. S2A), raising the intriguing possibility that this binding event might instead modulate other enolase activities such as one of the reported moonlighting functions (*11, 12*).

**Figure 3.**
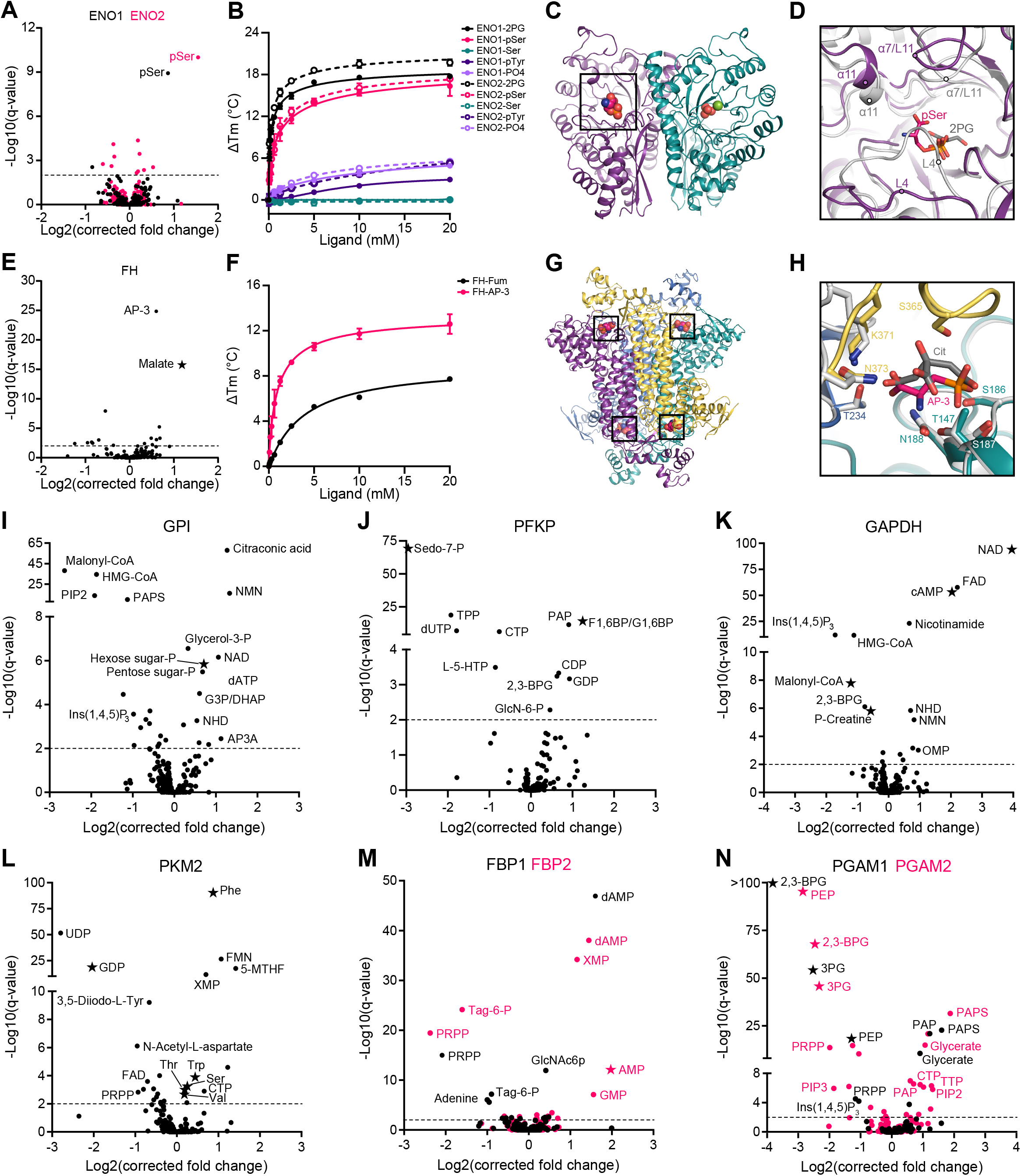
MIDAS reveals known and novel metabolite interactions with enzymes from human central carbon metabolism. **(A)** Metabolite interactions with enolase 1 (ENO1, black) and enolase 2 (ENO2, pink). **(B)** Ligand-induced DSF melting point analysis of ENO1 (solid lines, solid circles) and ENO2 (dotted lines, open circles) with 2-phosphoglycerate (2PG, black), phosphoserine (pSer, pink), serine (Ser, teal), phosphotyrosine (pTyr, purple), and phosphate (PO4, light purple). **(C)** X-ray crystal structure of the pSer-ENO2 complex (PDB 7MBH). pSer (black box), phosphate ion (orange and red spheres), magnesium ion (green sphere), monomers within the ENO2 dimer (purple and teal). **(D)** Magnified view of the ENO2 active site with pSer (pink) or 2-phoshoglycerate (2PG) (grey) bound (2PG-ENO2, PDB 3UCC) (*49*). Secondary structure labeled in the pSer-ENO2 (purple) and 2PG-ENO2 (light grey) structures. **(E)** Metabolite interactions with fumarase (FH). **(F)** Ligand-induced DSF melting point analysis of FH with fumarate (Fum, black) and 2-Amino-3-phosphonopropionic acid (AP-3, pink). **(B and F)** Line of best fit was determined using the specific binding and Hill slope equation from Prism 9. **(G)** X-ray crystal structure of the AP-3-FH complex (PDB 7LUB). AP-3 (black boxes), monomers within the FH tetramer (purple, yellow, teal, and light blue). **(H)** Magnified view of the FH active site with AP-3 (pink) or citrate (Cit, grey) bound (*E. coli* Cit-FH structure, light grey, PDB 1FUO) (*13*). Sidechains that coordinate the AP-3 interaction with FH are labeled and colored according to FH monomer from (G). **(I)** Metabolite interactions with glucose-6-phosphate isomerase (GPI). **(J)** Metabolite interactions with 6-Phosphofructokinase, platelet type (PFKP). **(K)** Metabolite interactions with glyceraldehyde-3-phosphate dehydrogenase (GAPDH). **(L)** Metabolite interactions with pyruvate kinase M2 (PKM2). **(M)** Metabolite interactions with fructose-1,6-bisphosphatase 1 (FBP1, black) and fructose-1,6-bisphosphatase 2 (FBP2, pink). **(N)** Metabolite interactions with phosphoglycerate mutase 1 (PGAM1, black) and phosphoglycerate mutase 2 (PGAM2, pink). **(A, E, I – N)** Volcano plots generated from MIDAS PMI data. Specific metabolites are labeled. Stars indicate a previously known human PMI primarily sourced from BRENDA (https://www.brenda-enzymes.org/index.php). Significant protein-metabolite interactions have a q-value < 0.01 (dotted line).

MIDAS identified 2-amino-3-phosphonopropionic acid (AP-3), a component of phosphonate metabolism (KEGG), as a putative interactor with fumarase, an enzyme in the tricarboxylic acid cycle that catalyzes the reversible hydration of fumarate to malate, which was also a significant hit (Fig. 3E). AP-3 induced the thermal stabilization of fumarase (Kd_app_ = 0.98 mM) with similar potency to its substrate, fumarate (Kd_app_ = 3.87 mM) (Fig. 3F). Kinetic assays demonstrated that AP-3 competitively inhibited fumarase (Fig. S2B), and, consistent with this, the crystal structure of the complex revealed that AP-3 binds in the active site of fumarase similarly to the known inhibitor citrate (Fig. 3G–H) (*13*). Although the consequences of fumarase modulation by AP-3 *in vivo* are unclear, these findings demonstrate that MIDAS can identify novel and functional protein-metabolite interations.

MIDAS datasets from additional proteins further confirmed the ability of MIDAS to identify known interactions with substrates, products, and regulators: Glucose-6-phosphate isomerase (GPI) with its substrates, glucose-6-phosphate and fructose-6-phosphate (hexose-6-phosphates) (Fig. 3I); phosphofructokinase (PFKP) with its product (F-1,6BP/G-1,6BP) and a putative alternative substrate, sedoheptulose-7-phosphate, (*14*) which is an intermediate in the pentose-phosphate pathway (Fig. 3J); glyceraldehyde-3-phosphate dehydrogenase (GAPDH) with its substrate (NAD),and regulators (cyclicAMP, creatine-phosphate, and malonyl-CoA) (*15–17*) (Fig. 3K); and the M2 isoform of pyruvate kinase (PK-M2) with GDP and multiple amino acid regulators (*18*) (Fig. 3L). In every case, MIDAS also highlighted intriguing previously unknown interactions with distinct and non-overlapping sets of metabolites. For example, acyl-CoAs, inositol phosphates, nicotinamides, adenine nucleotides, and downstream glycolytic intermediates were found to interact with GPI (Fig. 3I). We also observed interactions between PFKP and several di- and triphosphate nucleotides, thiamine pyrophosphate, as well as with L-5-hydroxytryptophan, an intermediate in the conversion of tryptophan to serotonin (Fig. J). Ins(3,4,5)P3, 2,3-BPG, and HMG-CoA were identified as novel binding partners for GAPDH (Fig. 3K), while PK-M2 interacted with flavins, a folate, and a thyroid hormone intermediate (Fig. 3L). PK-M2 was previously reported to be allosterically regulated by thyroid hormone T3, but the significance of this binding is unknown (*19*).

A comparison of MIDAS analyses for multiple isoforms of metabolic enzymes demonstrated both shared and unique metabolite interactions. Fructose bisphosphatase (FBP) catalyzes the conversion of fructose-1,6-bisphosphate to fructose-6-phosphate, a rate-limiting step in gluconeogenesis. Both isoforms (FBP1 and FBP2) interact with the known inhibitor AMP in addition to other nucleotide monophosphates. However, only FBP1 showed an interaction with glucosamine-6-phosphate, the rate-limiting intermediate in the hexosamine pathway, which emerges from fructose-6-phosphate (Fig. 3M). Similarly, isoforms of phosphoglycerate mutase (PGAM1 and PGAM2) interacted with a large set of metabolites, almost all of which were identical between them, with the exception of the soluble inositol phosphate inositol-1,3,4-P3 (PGAM1) and PIP2 and PIP3 (PGAM2) (Fig. 3N). This might reflect differential membrane recruitment and/or regulation of PGAM isoforms by phosphoinositide kinases, which are activated by growth factor signaling. These PMI data illustrate the power of MIDAS to enable the generation of hypotheses about potential novel regulatory events.

Lactate dehydrogenase (LDH) reversibly catalyzes the reduction of pyruvate to lactate coincident with the oxidation of NADH to NAD, a pivotal branchpoint in carbohydrate metabolism. Consumption of pyruvate by LDH competes with the mitochondrial uptake and oxidation by the TCA cycle to maximize ATP production. When mitochondrial pyruvate oxidation is limited, such as in hypoxia or aerobic glycolysis, LDH is critically important to regenerate NAD to enable continued glycolytic flux. Importantly, the LDH reaction is reversible and is also required to utilize lactate, a major circulating carbohydrate in mammals, as a fuel to support cellular functions (*20*). This firmly places LDH as a key node in carbohydrate metabolism.

MIDAS analysis of the two major isoforms of LDH, LDHA and LDHB, revealed interactions with several metabolites, most of which were common to both proteins (Fig. 4A). These included the cofactors NADH and NAD and the structurally related nucleotides nicotinamide mononucleotide (NMN) and FAD, as well as the competitive inhibitor, oxaloacetate (*21*), and other keto-acids related to the LDH substrates lactate and pryuvate. We also observed two other classes of interacting metabolites, adenosine nucleotides and free and acylated coenzyme A (Fig. 4A, B). To determine if either of these classes represent bona fide PMIs, we first used a thermal shift assay to measure the binding affinity of the three major adenosine nucleotides, AMP, ADP, and ATP, for LDHA and LDHB, and compared to the cofactor NAD (Fig. 4C). ATP interacted with both isoforms with a Kd_app_ = 0.636 mM and 0.697 mM, respectively, which is a biologically relevant affinity, given that the generally accepted intracellular steady state ATP concentration is 1–5 mM. The interactions of either LDH isoform with ADP and AMP may not be physiologically relevant given the disparity between the Kd values for each interaction and the cellular concentrations of ADP and AMP (~0.4 mM and ~0.04 mM, respectively (*22*) (Fig. 4C). Enzymatic activity assays of the two LDH isoforms further supported this conclusion as both AMP and ADP inhibited LDHA and LDHB only at supraphysiological concentrations (Fig. 4D). Interestingly, despite similar binding affinities to both LDHA and LDHB (Fig. 4D), ATP selectively inhibited only the LDHA isoform, with an IC_50_ of 2.3 mM. This discrepancy could potentially relate to the opposing effects of ATP binding on the thermal stability of the two proteins (Fig. 4C). The inhibition of LDHA by ATP appears to be competitive with NAD and lactate (Fig. S3A).

**Figure 4.**
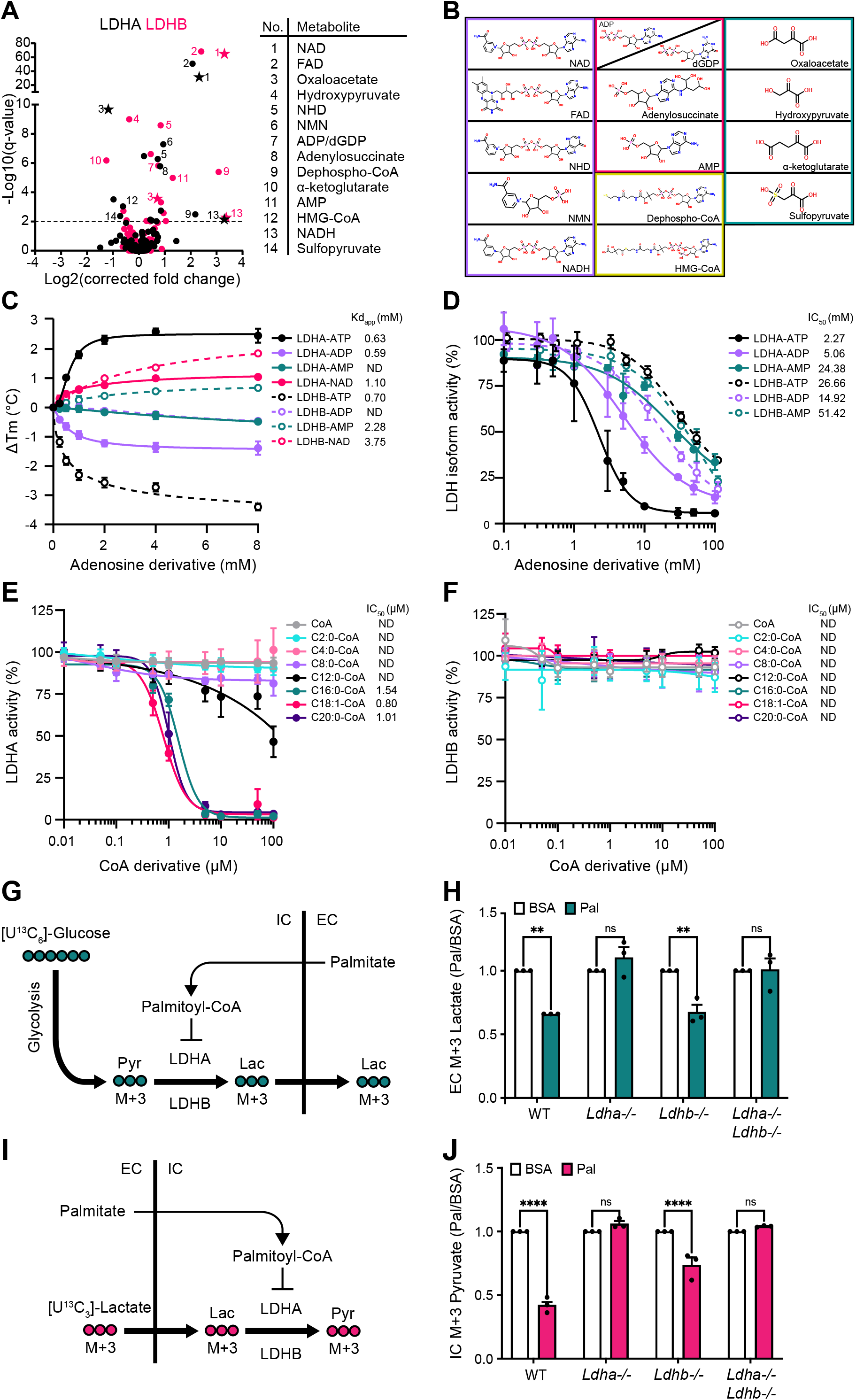
ATP and long-chain acyl-CoAs inhibit lactate dehydrogenase in an isoform-specific manner. **(A)** Metabolite interactions with lactate dehydrogenase A (LDHA, black) and lactate dehydrogenase B (LDHB, pink). Volcano plots generated from MIDAS PMI data. Specific metabolites are numbered and labeled. Stars indicate a previously known human PMI primarily sourced from BRENDA (https://www.brenda-enzymes.org/index.php). Significant protein-metabolite interactions have a q-value < 0.01 (dotted line). **(B)** Metabolite classes that interact with LDHA and LDHB from (A) (nicotinamides and dinucleotides, purple; adenosine nucleotide derivatives, pink; coenzyme A derivatives, yellow; keto acids, teal). **(C)** Ligand-induced DSF melting point analysis of LDHA (solid lines, filled circles) and LDHB (dotted lines, open circles) with adenosine triphosphate (ATP, black), adenosine diphosphate (ADP, light purple), adenosine monophosphate (AMP, teal), and nicotinamide adenine dinucleotide (NAD, pink). Apparent dissociation constant (Kd_app_) was determined from triplicate experiments using the specific binding and Hill slope equation from Prism 9. Mean ± SD is plotted from triplicate experiments. **(D)** Enzyme activity of LDHA (solid lines, filled circles) and LDHB (dotted lines, open circles) treated with ATP (black), ADP (light purple), or AMP (teal). **(E and F)** Enzyme activity of LDHA or LDHB treated with coenzyme A (CoA, grey), acetyl-CoA (C2:0-CoA, cyan), butyryl-CoA (C4:0-CoA, light pink), octanoyl-CoA (C8:0-CoA, light purple), lauroyl-CoA (C12:0-CoA, black), palmitoyl-CoA (C16:0-CoA, teal), oleoyl-CoA (C18:1-CoA, pink), and saturated arachidonoyl-CoA (C20:0-CoA, purple). **(D – F)** Half maximal inhibitory concentration (IC_50_) was determined from triplicate experiments using Prism 9; ND, not determined. Mean ± SD is plotted from triplicate experiments. **(G)** Schematic of [U^13^C_6_]-glucose metabolism in cells treated with palmitate-conjugated BSA. Pyruvate, Pyr; lactate, Lac; IC, intracellular; EC, extracellular. **(H)** Fold change of extracellular [U^13^C_3_]-lactate collected from the growth media of the indicated H9c2 cell lines in response to treatment with palmitate-conjugated BSA (Pal) relative to BSA control. **(I)** Schematic of [U^13^C_3_]-lactate metabolism in cells treated with palmitate-conjugated BSA. Pyruvate, Pyr; lactate, Lac; IC, intracellular; EC, extracellular. **(J)** Fold change of intracellular [U^13^C_3_]-pyruvate in indicated H9c2 cell lines in response to treatment with palmitate-conjugated BSA (Pal) relative to BSA control. **(H and J)** Experiments were performed in triplicate and mean ± SD is displayed. A students t-test was performed between Pal and BSA samples (p < 0.005, **; p < 0.00005, ****.

Next, we investigated the putative interaction between the LDH isoforms and coenzyme A (CoA) or CoA conjugated to short, medium, or long-chain fatty acids (i.e, acyl-CoAs). Esterification of long-chain (>12 carbons) fatty acids to CoA is required for their intracellular diffusion and transport into the mitochondrial matrix where they undergo β-oxidation to produce ATP. The accumulation of these species has been previously demonstrated to be a signal of carbon fuel excess (*23*). We observed that acyl-CoAs inhibited LDHA as a function of fatty acid chain length. Neither CoA alone nor any acyl-CoA with a fatty acid chain-length up to eight carbons affected enzyme activity, and C12:0-CoA only weakly inhibited LDHA, with an IC_50_ >100 μM (Fig. 4E). However, long-chain acyl-CoAs–C16:0-CoA (palmitoyl-CoA), C18:1-CoA (oleoyl-CoA) and C20:0-CoA (arachidoyl-CoA), none of which are in the MIDAS library–were all potent inhibitors of LDHA, with IC_50_ values of ~1 μM (Fig. 4E). The inhibition of LDHA by palmitoyl-CoA is non-competitive with respect to both NAD and lactate, suggesting that it is binding to LDHA outside of the active site (Fig. S3B). Intriguingly, LDHB, which shares 85% amino acid sequence identity with LDHA, was completely impervious to all tested acyl-CoAs, even at concentrations up to 100 μM (Fig. 4F).

Having observed that palmitoyl-CoA inhibited LDHA, but not LDHB, we employed two orthogonal approaches to test for a physical interaction. Using a thermal stability assay, we found that low micromolar concentrations of palmitoyl-CoA, similar to the IC_50_, induced the formation of a distinct thermo-labile species of LDHA, while inducing a thermo-stable species of LDHB (Fig. S3C). These data indicate that LDHA and LDHB both directly interact with palmitoyl-CoA with a physiologically relevant low micromolar affinity. Next, we found that purified LDHA and LDHB bound to palmitoyl-CoA immobilized on agarose beads, and that the binding for either protein was disrupted by elution with palmitoyl-CoA but not with buffer or acetyl-CoA (C2:0-CoA) (Fig. S3D).

Given that palmitoyl-CoA inhibited LDHA at concentrations within the physiological range, we next tested whether this inhibition is relevant in intact cells. We performed metabolic tracing experiments using H9c2 rat cardiomyoblasts in which we knocked out *Ldha*, *Ldhb*, or both (Fig. S3E). We treated cells with ^13^C-labeled glucose in the presence or absence of BSA-palmitate, which allows for efficient delivery of the fatty acid into the cell where it can esterified to palmitoyl-CoA (Fig. 4G). We then used mass spectrometry to measure the uptake and assimilation of ^13^C into lactate. All four cell lines (WT, *Ldha*−/−, *Ldhb*−/− and *Ldha*−/− *Ldhb*−/−) showed a similar (~80%) increase in intracellular palmitate following incubation with its BSA-conjugate (Fig. S3F). Importantly, palmitate decreased the labelling of lactate from ^13^C-glucose, but only in wild-type (WT) and *Ldhb*−/− cells i.e., in cells in which LDHA is still present (Fig. 4H, S3G–H), implying that palmitate inhibition of glucose-to-lactate conversion is completely dependent upon LDHA in this cell line. This result indicates that one or more steps between glucose uptake and LDHA-catalyzed lactate synthesis is inhibited by palmitate treatment. To probe more specifically the uptake and LDH-mediated oxidation of lactate to pyruvate, we performed analogous experiments with ^13^C-lactate (Fig. 4I). Again, we found that treatment with palmitate inhibited the generation of ^13^C-pyruvate in WT and LDHB−/− cells, but pyruvate labeling in *Ldha*−/− or *Ldha*−/− *Ldhb*−/− cells was unaffected (Fig. 4J, S3I–J). These data suggest a novel mode of regulatory crosstalk between fatty acid and carbohydrate metabolism.

It is intriguing that we found that both ATP and long-chain acyl-CoAs preferentially inhibit LDHA, but not LDHB. LDHA and LDHB, the two dominant isoforms of lactate dehydrogenase, are expressed in a tissue-specific pattern such that the liver almost exclusively expresses LDHA, while the heart has high expression of LDHB (Fig. S4A, B). Importantly, the IC_50_ for inhibition by ATP is easily within the range of normal intracellular ATP concentrations, suggesting that LDHA might be partially inhibited in all cells with normal energy status. Given that the liver, the most LDHA-dominant tissue, is capable of catabolizing multiple substrates, inhibition by ATP might be a mechanism to spare carbohydrates for those cell types that are dependent upon them. This is particularly important given the recent demonstration that lactate may be a major carbohydrate fuel consumed by some tissues (*20*). Likewise, our observation that long-chain acyl-CoAs inhibit LDHA could explain a previously described physiological phenomenon wherein fatty acids, released from the adipose tissue during fasting, inhibit lactate production and increase glucose production in the liver (*24*). We hypothesize that fatty acyl-CoA-mediated inhibition of LDHA would redirect pyruvate toward gluconeogensis and away from excretion following its conversion to lactate. Another potential implication of these results lies in the substantial interest in LDHA-specific inhibitors to block aerobic glycolysis in cancers (*25, 26*). Perhaps the mechanism(s) employed by ATP and acyl-CoAs could be exploited therapeutically.

This fatty acid-carbohydrate inter-pathway metabolic regulation is just one potential example of the myriad metabolite-level regulatory events that enforce organismal homeostasis, which is vital to appropriately respond to stressors such as the feed-fast cycle, exercise, and infection. We propose that interactions between proteins and metabolites mediate much of this control. We have validated MIDAS as a robust platform for the discovery of these critical mechanisms, particularly for the detection of very low affinity interactions, which include most of those involving high abundance cellular metabolites. In addition to recent discoveries of functionally important PMIs (*27–29*), MIDAS now identified hundreds of known and novel putative interactions with the enzymes of central carbon metabolism. Together, these serve as a roadmap for identifying new modes of metabolic regulation as well as previously undescribed alternative substrates. We propose that the comprehensive discovery of such interactions will revolutionize our understanding of cell biology and that MIDAS can empower a renewed focus on this challenging and critical important area of biology.

## Supporting information

Data S1. MIDAS metabolite library

Data S2. FIA-MS properties of MIDAS metabolites

Data S3. MIDAS proteins

Data S4. MIDAS protein-metabolite interactions

## Acknowledgements

We thank members of the Rutter lab for helpful discussions and comments on the manuscript. We thank Roche and Navitor for providing proteins for MIDAS analyses. We thank the University of Utah Mutation Generation and Detection Core for providing CRISPR reagents, cell genotyping services and construction of bacterial expression vectors. We thank the University of Utah Drug Discovery Core Facility for generating the *Ldha* and *Ldhb* mutant cells. Metabolomics analysis was performed in the Metabolomics Core Facility at the University of Utah. Mass spectrometry equipment was obtained through NCRR Shared Instrumentation Grant 1S10OD016232-01, 1S10OD018210-01A1 and 1S10OD021505-01. This work was supported by T32DK091317 and T32DK007115 to KGH, U54DK110858, R35GM131854, and a grant from Calico Life Sciences, LLC to JR. JR is an investigator of the Howard Hughes Medical Institute. KGH and JR are inventors of MIDAS technology that has been licensed to Atavistik Bio, for which KGH is a consultant and JR is a founder.

## Materials and Methods

### MIDAS metabolite library construction and storage

The MIDAS metabolite library (Fig. S1A) was constructed by extracting and cross-referencing primary and secondary metabolites from KEGG and HMDB, with a focus on endogenous and exogenous compounds that were quantified, detected, or predicted in human metabolism. All metabolites used in this study were purchased from Sigma-Aldrich, Cayman Chemicals, Avanti Polar Lipids, Enamine, Combi-Blocks, Inc, or custom sourced using Aldrich Market Select (Data S1). Metabolites were solvated to 10 mM in molecular grade water (Sigma-Aldrich W4502) or DMSO (Sigma-Aldrich D1435) and, where necessary to increase solubility, titrated with acid or base. The MIDAS metabolite library was arrayed 1 mL per well in 96-deep well storage plates (Greiner 780280), sealed with aluminum foil seals (VWR 60941-112), and stored at −80°C. When working stocks were needed, metabolites were moved from the deep well storage plates and arrayed, 50 μL per well, across multiple, single-use 384-well small volume plates storage plates (Greiner 781280), sealed with aluminum foil seals (VWR 60941-112), and stored at −80°C. Metabolite library management and manipulation was conducted on a Beckman Coulter Biomek NX^p^ SPAN-8 liquid handling robot.

### MIDAS metabolite library validation and pooling

Metabolite accurate mass, adduct, ionization, and detection parameters were determined using a flow injection analysis mass spectrometry (FIA-MS) scouting approach to design four defined metabolite screening pools (Fig. S1B) (Data S2). Briefly, 20 pmol of each metabolite from the MIDAS metabolite library was independently assayed in positive and negative mode in technical quadruplicate 1 μL injections with interspersed blank injections by FIA-MS on a binary pump Agilent 1290 Infinity UHPLC system operated with a flow rate of 0.1 mL/min coupled to an Agilent 6550 ESI-QTOF MS. The following mobile phases were used for FIA-MS scouting: 20 mM formic acid pH 3 (Sigma-Aldrich F0507), 10 mM ammonium acetate pH 5 (Sigma-Aldrich 73594), 10 mM ammonium acetate pH 6.8 (Sigma-Aldrich 73594), and 10 mM ammonium bicarbonate pH 9 (Sigma-Aldrich 09830). Source conditions consisted of 250°C gas temp, 11 L/min gas flow, 20 psig nebulizer, 400° sheath gas temperature, 12 L/min sheath gas flow, and 2000 V nozzle voltage. Agilent MassHunter 7 software was used to qualitatively validate and quantify metabolites. The optimal signal for each metabolite was determined by integrating the area under the curve of the extracted ion chromatogram (XIC-AUC) for each metabolite adduct at the various mobile phase pH and instrument polarity. The optimal adduct, pH, polarity, metabolite solvent, and, if necessary, isomer family of each metabolite was considered to construct four unique and defined MIDAS metabolite screening pools (Data S1).

### MIDAS protein-metabolite screening

The day of MIDAS screening, a number of MIDAS metabolite library, 384-well small volume working stock plates, corresponding to the number of proteins to be screened (eight proteins per plate), were defrosted at 30°C for 5 minutes and metabolites were combined *de novo* to generate four predetermined MIDAS screening pools (Data S1). The MIDAS screening pools were prepared in LC-MS grade 150 mM ammonium acetate pH 7.4 (Sigma-Aldrich 73594) and pH-adjusted with ammonium hydroxide (Sigma-Aldrich 338818). The majority of metabolites were prepared to a final screening concentration of 50 μM in the metabolites pools, with a subset at higher or lower concentration dependent on their FIA-MS ionization properties (Data S1). For each metabolite pool, 8 μL of target protein (Fig. S1C and Data S3) was arrayed in a minimum of a triplicate across a 10 kDa MWCO 96-well microdialysis plate (SWISSCI Diaplate™) and sealed with aluminum foil seals (Beckman Coulter 538619) to create the protein chambers. To the reverse side, 300 μL of metabolite pool was aliquoted per target protein replicate and sealed with aluminum foil seals (Beckman Coulter 538619) to create the metabolite chambers. Where necessary and just prior to screening, proteins provided in alternative buffer systems were *in situ*, sequentially exchanged into 150 mM ammonium acetate pH 7.4 (Sigma-Aldrich 73594) on the 96-well microdialysis screening plate (SWISSCI Diaplate™). Loaded dialysis plates were placed in the dark at 4°C on a rotating shaker (120 rpm) and incubated for 40 hours. Post-dialysis, protein and metabolite chamber dialysates were retrieved, sample volume normalized and diluted 1:10 in 80% methanol (Sigma-Aldrich 1060351000) to precipitate protein, incubated 30 mins on ice, and centrifuged at 3200 x g for 15 mins to sequester precipitated protein. Processed protein and metabolite chamber dialysates were retrieved and arrayed across a 384-well microvolume plate (Thermo Scientific AB-1056), sealed with a silicon slit septum cap mat (Thermo Scientific AB-1171), and placed at 4°C for FIA-MS analysis.

### MIDAS flow injection analysis mass spectrometry analysis

All MIDAS metabolite pool FIA-MS was performed on a Shimadzu Nexera HPLC system equipped with binary LC-20AD_XR_ pumps and a SIL-20AC_XR_ autosampler coupled to a SCIEX X500R ESI-QTOF MS. Briefly, 2 μL of each processed protein and metabolite chamber dialysate (~10 pmoles per metabolite, depending on metabolite) was injected in technical triplicate with blanks injections interspersed between technical replicates. Mobile phase flow rate was 0.2 mL/min. The following mobile phases were used according to the MIDAS metabolite pool being analyzed: pool 1, 5 mM ammonium acetate pH 5 (Sigma-Aldrich 73594), 50% methanol (Honeywell LC230-4); pools 2 and 4, 5 mM ammonium acetate pH 6.8 (Sigma-Aldrich 73594), 50% methanol (Honeywell LC230-4); pool 3, 10 mM formic acid pH 3 (Sigma-Aldrich F0507), 50% methanol (Honeywell LC230-4). Pools 1 and 2 were analyzed in positive mode and pools 3 and 4 were analyzed in negative mode. Source conditions consisted of 40 psi for ion source gas 1 and 2, 30 psi curtain gas, 600°C source temperature, and +5500 V or −4500 V spray. Method duration was 1 min. All target proteins for a given metabolite pool and MS method were analyzed together before switching FIA-MS methods. Between FIA-MS methods, the Shimadzu Nexera HPLC system and SCIEX X500R ESI-QTOF MS where equilibrated for 40 min to the next FIA-MS method. Auto-calibration of positive or negative mode was performed approximately every 45 mins at the beginning of a target protein-metabolite pool batch to control detector drift. Non-dialyzed MIDAS metabolite pools were assayed at the beginning, middle, and end of each metabolite pool method batch to control detector sensitivity.

### MIDAS data processing and analysis

MIDAS FIA-MS spectra were processed in SCIEX OS 1.6 software using a targeted method to determine metabolite abundances in the protein chamber and metabolite chamber by integrating the mean area under the curve for each extracted ion chromatogram. If necessary, up to one dialysis replicate per pool per protein was removed if processing or autosampling abnormalities were identified. For each dialysis replicate, log2(fold change) for each metabolite was calculated as the difference between the log2 abundance in the protein chamber and metabolite chamber. Log_2_(fold change) for metabolite isomers (e.g. L-Leu/L-Ile/L-Allo-Ile) within the same screening pool were collapsed to a single entry prior to further data processing leading to 333 unique metabolite isomer analytes. Using the replicate protein-metabolite log2(fold change) values as input, a processing method was developed in R (https://github.com/KevinGHicks/MIDAS) to capture and remove extreme outliers and non-specific systematic variation and to determine significant protein-metabolite interactions. Briefly, for each dialysis replicate set, up to one outlier was removed using a z-score cutoff of five (<0.2% of observations). Technical replicates were then averaged yielding one fold-change summary per protein-metabolite pair. To remove fold-change variation that was not specific to a given protein-metabolite pair, the first three principal components of the total screening dataset were removed on a per metabolite pool basis by subtracting the projection of the first three principal components, creating log_2_(corrected fold change). Protein-metabolite z-scores were determined by comparing the target protein-metabolite log_2_(corrected fold change) to a no-signal model for that metabolite using measures of the central tendency (median) and standard deviation (extrapolated from the inter quartile), which are robust to the signals in the tails of a metabolite’s fold-change distribution. Z-scores were false-discovery rate controlled using Storey’s q-value (*30*) and protein-metabolite interactions with q-values < 0.01 were considered significant. Since correcting for non-specific binding, and estimating metabolite-specific standard deviation both benefit from the inclusion of additional proteins, MIDAS data from 122 anonymized proteins were analyzed alongside the 38 proteins specifically considered in this study. The complete MIDAS protein-metabolite interaction dataset for mTORC1 regulators and enzymes of central carbon metabolism can be found in Data S4.

### MIDAS proteins

All presented proteins analyzed by MIDAS were prepared and provided by collaborators (Data S3) using common protein expression and purification techniques. Proteins were received snap frozen on dry ice from outside sources or on wet ice from local sources. Prior to MIDAS screening, protein quality was assessed by 12.5% SDS-PAGE and concentration was determined by A280 on a NanoDrop One UV-Vis spectrophotometer using the molecular weight and calculated extinction coefficient (M^−1^ · cm^−1^) of each protein construct. Proteins were screened by MIDAS at the concentrations indicated in Data S3.

### Electrum

MIDAS protein-metabolite interaction data for enzymes of central carbon metabolism were visualized for intra- and inter-pathway relationships using Electrum (v0.0.0; https://github.com/Electrum-app/Electrum), with q-value cutoff < 0.01, and the 1-D scaling option enabled.

### Differential scanning fluorimetry

Thermal differential scanning fluorimetry (DSF) was performed similar to Niesen *et al* (*31*). Briefly, DSF thermal shift assays were developed to assess protein melting point (Tm) and thermal stability in the presence of putative small molecule ligands: 2-phosphoglycerate (2PG, Sigma-Aldrich 73885), phosphoserine (pSer, Sigma-Aldrich P0878), phosphotyrosine (pTyr, Sigma-Aldrich P9405), phosphate (PO4, Acros Organics 424395000), fumarate (Fum, Sigma-Aldrich 47910), 2-amino-3-phosphonopropionic acid (AP-3, Sigma-Aldrich A4910), ATP (Sigma-Aldrich A2383), ADP (Sigma-Aldrich 01905), AMP (Sigma-Aldrich A2252), NAD (Sigma-Aldrich N1636), and palmitoyl-CoA (C16:0-CoA, Avanti 870716). Where indicated, DSF experiments were performed using either the standard SYPRO orange fluorescent system or PROTEOSTAT® Thermal shift stability assay kit (ENZO 51027). A final concentration reaction mixture of 10 μL containing 25mM HEPES pH 7.4, 50mM NaCl, 0.1 mg/mL (SYPRO system) or 0.75 mg/mL (PROTEOSTAT system) target protein, 7.5X SYPRO orange (Sigma-Aldrich S5692) or 1x PROTEOSTAT® reagent, and the indicated concentration of putative ligand was arrayed across a MicroAmp™ optical 384-Well reaction plate (Thermo Scientific 4309849) and sealed with MicroAmp™ optical adhesive film (Thermo Scientific 4360954). Protein denaturation was measured in sextuplicate technical replicates for SYPRO orange and PROTEOSTAT experiments with an excitation of 470 nm and emission of 580 nm on an Applied Biosystems Quantstudio 7 Flex from 25°C to 95°C at a ramp rate of 0.05°C/second. DSF experiments were performed in triplicate. Protein Thermal Shift software 1.4 (Applied Biosystems) was used to interpret and determine protein Tm from the first derivative of the fluorescence emission as a function of temperature (dF/dT). A change in ligand-induced protein melting point (ΔTm) was determined from the difference of the ligand induced Tm and no-ligand control Tm. Apparent binding affinity (Kd_app_) was determined by fitting the specific binding and Hill slope equation to ΔTm as a function of ligand concentration in GraphPad Prism 9 software.

### Fumarase competitive inhibition assay

The competitive inhibition of human fumarase activity in the presence of 2-amino-3-phosphonopropionic acid (AP-3, Sigma-Aldrich A4910) was fluorometrically assessed using a coupled enzyme assay. Briefly, the rate limiting hydration of fumarate to malate by fumarase provides substrate, malate, for excess malate dehydrogenase to generate oxalacetate and NADH. Fumarase reaction rate was assessed at room temperature in triplicate with a final reaction volume of 100 μL composed of 50 mM Tris-HCl pH 9.4, 61 μg/mL human fumarase, excess porcine heart malate dehydrogenase (Sigma-Aldrich 442610-M), 1 mM NAD (Sigma-Aldrich N1636), and varying concentrations of fumarate (0 – 40 mM, Sigma-Aldrich 47910) and AP-3 (0 – 10 mM, Sigma-Aldrich A4910) (Fig. S2). Fumarate and AP-3 were added simultaneously to initiate the reaction. The production of NADH was quantified fluorometrically in a black, clear bottomed 96-well plate (Sigma-Aldrich CLS3603) on a Biotek Synergy Neo plate reader with 360 nm excitation and 460 nm emission over 10 minutes and fumarase reaction rate was determined from the linear range of increasing NADH signal. A Lineweaver-Burke linear regression and non-linear regression competitive inhibition model of human fumarase between fumarate and AP-3 were fit using GraphPad Prism 9 software from triplicate competitive inhibition experiments.

### Enolase 2 Activity Assay

Human enolase 2 (ENO2) activity was measured in the presence of phosphoserine using a coupled enzyme kinetic assay similar to Satani *et al* (*32*). Briefly, enolase converts 2-phosphoglycerate (2PG) to phosphoenolpyruvate (PEP) and water. Substrate, 2PG, was provided near the measured Km. Excess pyruvate kinase (PK) / lactate dehydrogenase (LDH) enzymes from rabbit muscle (Sigma P0294), ADP, NADH were added to solution to ensure that dehydration of 2PG by enolase was the rate-limiting step. Enolase reaction rate was assessed at room temperature in triplicate with a final reaction volume of 100 uL composed of 50 mM HEPES pH 7.4, 0.5 mM MgCl2, 100 uM NaCl, 1.75 mM ADP, 200 uM NADH, 12.8 U PK, 18.4 U LDH, 500 ug ng of ENO2, 30 uM 2PG, with varying concentrations of phosphoserine (pSer). 2PG was used to initiate the coupled enzyme reaction, and the conversion of NADH to NAD by LDH was quantified fluorometrically in a black, clear bottomed 96-well plate (Sigma-Aldrich CLS3603) on a Biotek Synergy Neo plate reader with 360 nm excitation and 460 nm emission over 10 minutes. Enolase reaction rate was determined for the linear range of decreasing NADH signal. IC50 was determined using a sigmoidal, 4PL non-linear regression in Prism 9 from triplicate experiments.

### Enolase X-ray crystallography

Crystals of Human Enolase 2 in complex with the phosphoserine ligand were prepared via hanging drop vapor diffusion at 20 °C. 9 mg/ml- Human Enolase 2 protein solution with 2 mM phosphoserine was pre-incubated on ice for 10 min prior to being mixed in 1:1 ratio (protein:reservoir solution) with 100 mM Bis Tris, 200 mM ammonium acetate and 21% (w/v) PEG 3350 at pH 6.5. Orthorhombic crystals grew within 3 days and were subsequently cryoprotected with 100 mM Bis Tris, 200 mM ammonium acetate, 32% (w/v) PEG 3350 and 2 mM phosphoserine. X-ray diffraction data were collected at the Advanced Photon Source, synchrotron beamline 22-ID, equipped with Si(III) monochromator and EIGER CCD detector. The diffraction data was processed and integrated using iMOSFLM (*33*). POINTLESS (*34*) was used to identify the bravais lattice and space group and AIMLESS (*35*) was used for scaling. The phase information was obtained by molecular replacement using PHASER (*36*) with a homodimer of Human Enolase 2 (PDB 4ZCW) as the search model. Iterative cycles of manual model building and refinement were performed within Phenix (*37*) and COOT (*38*) software. Diffraction data and refinement statistics are summarized in Table S1.

### Fumarase X-ray crystallography

Human fumarase (HsFH) was produced and purified as previously described (*39*). The co-crystallization experiments were carried out by using the sitting drop method. Protein solution (4 mg/mL in 50 mM Tris-HCl (Sigma-Aldrich), pH 8.5, 150 mM KCl (J.T.Baker) was incubated with 20 mM of D-2-amino-3-phosphono-propionic acid (AP-3). 2 μL of protein solution was mixed with 2 μL of reservoir solution, and allowed to equilibrate against 500 μl of reservoir solution at 21°C. Crystals occur over the course of 3 days in drops where the reservoir contained 100 mM Hepes pH 7.5 (Sigma-Aldrich), 1% v/v 2-methylpentanediol (MPD) (Sigma-Aldrich) and 18% (w/v) PEG 10 K (Sigma-Aldrich) and 25% (v/v) glycerol. Prior to data collection, HsFH crystals were soaked in a cryoprotectant solution (100 mM Hepes pH 7.5, 1% v/v 2-methylpentanediol (MPD), and 18% m/v PEG 10 K, 25% v/v glycerol (Labsynth), harvested with cryo loops, and flash-cooled in liquid nitrogen. The data set was collected at 100 K on a synchrotron facility (MANACA beamline - SIRIUS, Brazil) using a PILATUS 2M detector (Dectris). 3600 frames with an oscillation step of 0.1° were collected using an exposure time of 0.1 s per image with a crystal-to-detector distance of 120.05 mm. The images of X-ray diffraction were processed with XDS (*40*) package, and the structure of HsFH was solved by molecular replacement implemented in Molrep (*41*) program, and using the human fumarase structure (PDB ID: 5UPP) (*39*) as a template. The structure was refined with Refmac5 (*42*) intercepted with manual map inspection and model building using Coot (*38*). The quality of the model was regularly checked using MolProbity (*43*). Diffraction data and refinement statistics are summarized in Table S2. The refined atomic coordinates and structure factors were deposited in the PDB with the accession code 7LUB.

### Lactate dehydrogenase activity assay

Human lactate dehydrogenase A (LDHA) and lactate dehydrogenase B (LDHB) activity were assessed in the presence of putative nucleotide and fatty acyl-CoA derivatives using a standard NADH fluorometric assay. Briefly, lactate dehydrogenase reversibly converts lactate and NAD to pyruvate and NADH. With the exception of the competitive inhibition assay, LDHA and LDHB activity assays were operated near their measured substrate and cofactor Km. Lactate dehydrogenase reaction rate was assessed at room temperature in triplicate with a final reaction volume of 100 μL composed of 75 mM Tris pH 7.4, 67.2 ng/ml LDHA or 75 ng/mL LDHB, 6.5 mM Lactate (Sigma-Aldrich L6402) and 200 μM NAD (Sigma-Aldrich N1636) for LDHA and 1mM lactate and 1.25 mM NAD for LDHB, with varying concentrations of putative ligand, as indicated: ATP (Sigma-Aldrich A2383), ADP (Sigma-Aldrich 01905), AMP (Sigma-Aldrich A2252), CoA (Avanti 870701), C2:0-CoA (Avanti 870702), C4:0-CoA (Avanti 870704), C8:0-CoA (Avanti 870708), C12:0-CoA (Avanti 870712), C16:0-CoA (Avanti 870716), C18:1-CoA (Avanti 870719), and C20:0-CoA (Avanti 870720). For competitive inhibition assay, the concentrations of lactate or NAD were varied, accordingly. In all circumstances, NAD was used to initiate the lactate dehydrogenase reaction. The production of NADH was quantified fluorometrically in a black, clear bottomed 96-well plate (Sigma-Aldrich CLS3603) on a Biotek Synergy Neo plate reader with 360 nm excitation and 460 nm emission over 10 minutes and lactate dehydrogenase reaction rate was determined from the linear range of increasing NADH signal. IC_50_ were determined using a sigmoidal, 4PL non-linear regression in Prism 9 from triplicate experiments. Non-linear regression competitive or non-competitive inhibition modeling of LDHA between lactate or NAD and ATP or palmitoyl-CoA were fit using GraphPad Prism 9 software from triplicate experiments.

### Palmitoyl-CoA-Agarose pull-down assay

LDHA and LDHB interaction with palmitoyl-CoA was assessed using a pull-down, competitive elution assay. Briefly, 30 μL per pull-down of palmitoyl-CoA conjugated agarose beads (Sigma-Aldrich 5297) were buffer exchanged into pull-down buffer (75mM Tris HCl pH 7.4). In a final volume of 300 μL, 0.2 mg/mL of LDHA or LDHB protein were combined with buffer exchanged palmitoyl-CoA agarose beads, a loading control was saved, and the mixture was incubated overnight at 4°C with gentle agitation. Post-incubation, pull-down reactions were washed 5x in 100 μL of pull-down buffer and the final wash was saved for analysis. Following the fifth wash, 100 μM of acetyl-CoA or palmitoyl-CoA or equivalent volume of pull-down buffer were added to the reactions and incubated overnight at 4°C. In the morning, the pull-down reactions were centrifuged to pellet beads, the supernatant was collected and concentrated as the eluate fraction, the beads were collected as the bound fraction, and all samples were boiled for 5 minutes in 4x Laemmli sample buffer and analyzed by SDS-PAGE for presence of LDHA or LDHB.

### Tissue culture

H9c2 myoblastic cell line (ATCC CRL-1446) was purchased from ATCC and routinely maintained in DMEM media supplemented with 10% FBS and 1% PenStrep in 5% CO2 and 37°C.

### *Ldha* and *Ldhb* mutant cell lines

*Ldha* and *Ldhb* knock out H9c2 cell lines were generated using CRISPR-Cas9 to excise the first coding exon of each gene. Single guide modified synthetic sgRNAs were obtained from Synthego and Hifi-Cas9 was obtained from IDT (cat# 1081060). Pairs of ribonucleoprotein (RNP) complexes targeting upstream and downstream of the first coding exon for each gene were co-electroporated using a Lonza 4D Nucleofector system (https://knowledge.lonza.com/cell?id=1016&search=H9c2). The N20 sgRNA target sequences used were GAGTGCAACGCTCAACGCCA and TCCACAGGCTTGTGACATAA for *Ldha* and TCCATGCATGTAAAGCACAT and AAGACAGCACAACTCTATAG for *Ldhb*. Off-targets for these sgRNAs were screened using CasOT (*44*). Nucleofected cells were plated as single clones and clones were screened for the expected genomic deletion and presence of the WT allele using PCR and these results were confirmed using Western blotting.

### Cell lysate and western blotting

Harvested cells were washed with PBS and lysed in RIPA buffer (50 mM Tris, 150 mM NaCl, 0.1% SDS, 0.5% sodium deoxycholate, 1% NP-40) supplemented with protease and phosphatase inhibitors. Protein concentration was quantified with the Pierce BCA Protein Assay Kit. Samples were mixed with 4x sample loading buffer and incubated for 5 min at 95°C. 30 μg of total protein lysate was resolved on SDS polyacrylamide gel according to standard procedure at 20 mA per gel and blotted onto a nitrocellulose membrane 0.45 μm (GE Healthcare) via Mini Trans-blot module (Bio-Rad) at a constant voltage (100 V) for 2 h. After blocking with 5% non-fat milk (Serva)/Tris-buffered saline with 0.05% Tween 20 (TBS-T) for 1 h, the membrane was incubated overnight in 5% bovine serum albumin (Sigma-Aldrich), TBS-T with primary antibody against LDHA (Cell Signaling Technology 2012S, 1:1000), LDHB (Abcam ab240482, 0.1 g/ml), and GAPDH (Cell Signaling Technology 97166, 1:1000). Next day, the membrane was washed with TBS-T and incubated with corresponding fluorophore-conjugated secondary antibody (Rockland Immunochemical RL611-145-002, 1:10000) in 1% non-fat milk/TBS-T for 1 h. The membrane was then washed again with TBS-T and fluorescence was assessed with Odyssey CLx imaging system (LI-COR Biosciences).

### Metabolite extraction

The procedures for metabolite extraction from cultured cells are described in previous studies (*45–47*). Briefly, adherent cells were grown in 10 cm plates in biological triplicate to 80% confluence, medium was rapidly aspirated and cells were washed with cold 0.9% NaCl TC grade (Sigma-Aldrich S8776-100ML) on ice. 3 mL of extraction solvent, 80% (v/v) LC/MS grade methanol/water (Fisher Scientific W6-1, A456-1) cooled to −80°C, was added to each well, and the dishes were transferred to −80°C for 15 min. Cells were then scraped into the extraction solvent on dry ice. Additionally, 300mL of media was collected and processed from each sample pre and post experiment. All metabolite extracts were centrifuged at 20,000 x g at 4°C for 10 min. Each sample was transferred to a new 1.5 mL tube. Finally, the solvent in each sample was evaporated in a Speed Vacuum, and stored at −80°C until they were run on the mass spectrometer.

### [U-13C6]-Glucose and [U-13C3]-Lactate labeling with or without palmitate

Cells were grown to 80% confluence in 10 cm plates with standard culture medium at which point 10 μM of the MPC inhibitor UK5099 (Sigma-Aldrich PZ0160-5MG) was added for 48 hours to facilitate lactate production. Cells were subsequently washed with sterile PBS and either free BSA or BSA conjugated to palmitate (Caymen Chemical 29558) was added to culture media containing either [U-13C6]-L-glucose, or [U-13C3]-L-lactate (Cambridge Isotope Laboratories CLM-1396, CLM-1579-PK), supplemented with dialyzed Fetal Bovine Serum (Thermo Scientific A3882001) and incubated for 4 hours. Metabolites were extracted as described above. Data was corrected for naturally occurring 13C isotope abundance before analysis as described in Buescher *et al* (*48*). All data expressed as mean ± SD unless otherwise indicated. Student’s t test was used for 2 group comparison. One-Way ANOVA and Sidak’s comparisons were used for multigroup comparison. p < 0.05 were considered statistically significant. Statistical analyses and graphics were carried out with GraphPad Prism 9 software.

### Metabolomic analysis

The levels of metabolites in the H9c2 cells were measured by gas chromatography–mass spectroscopy (GC-MS) analysis. All GC-MS analysis was performed with a Waters GCT Premier mass spectrometer fitted with an Agilent 6890 gas chromatograph and a Gerstel MPS2 autosampler. Dried samples were suspended in 40 μL of a 40 mg/mL O-methoxylamine hydrochloride (MOX) in pyridine and incubated for 1 h at 30°C. 10 μL of N-methyl-N-trimethylsilyltrifluoracetamide (MSTFA) was added automatically via the autosampler and incubated for 60 min at 37°C with shaking. After incubation 3 μL of a fatty acid methyl ester standard solution was added via the autosampler. Then 1 μL of the prepared sample was injected to the gas chromatograph inlet in the split mode with the inlet temperature held at 250°C. A 10:1 split ratio was used for analysis. The gas chromatograph had an initial temperature of 95°C for one minute followed by a 40°C/min ramp to 110°C and a hold time of 2 min. This was followed by a second 5°C/min ramp to 250°C, a third ramp to 350°C, then a final hold time of 3 min. A 30 m Phenomex ZB5-5 MSi column with a 5 m long guard column was employed for chromatographic separation. Helium was used as the carrier gas at 1 mL/min. Data was extracted from each chromatogram as area under the curve for individual metabolites. Each sample was first normalized to the added standard d4-succinate to account for extraction efficiency followed by normalization to cell number. Due to this being a broad scope metabolomics analysis, no normalization for ionization efficiency or concentration standards was performed.

---

**Figure S1.**
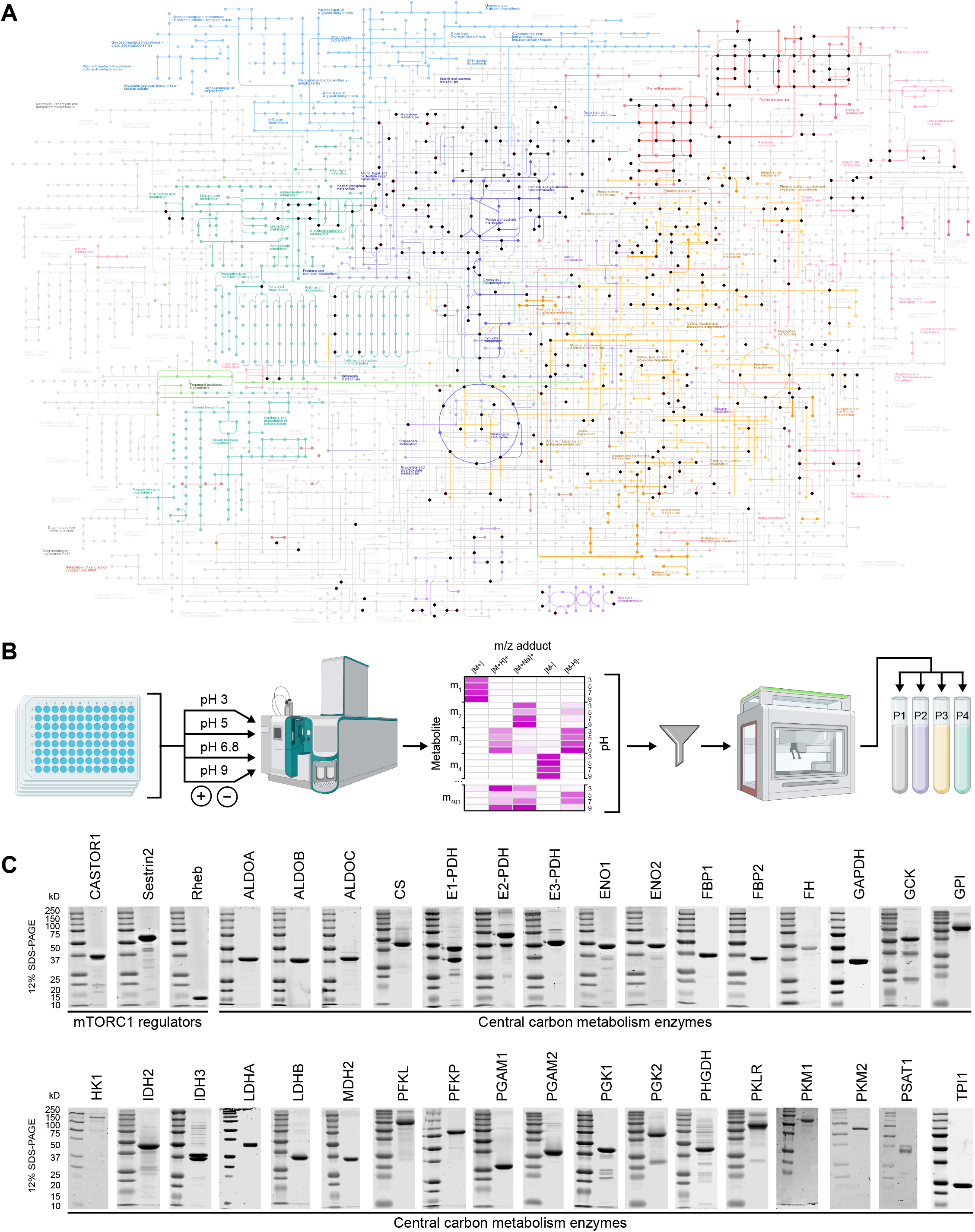
MIDAS metabolite library construction and validation and screened proteins. **(A)** The MIDAS metabolite library overlaid on KEGG human metabolic pathways (https://www.genome.jp/kegg/pathway/map/map01100.html). Human pathways, colored; non-human pathways, light grey. Metabolites from the MIDAS metabolite library, black spheres. **(B)** FIA-MS scouting method to determine optimal MIDAS metabolite pools. Metabolites from the MIDAS metabolite library were arrayed across five 96-well plates in water. Each metabolite was individual analyzed at mobile phase pH 3, 5, 6.8, and 9, in positive and negative mode, by FIA-MS. For each metabolite (m_1_ – m_401_) analyzed by FIA-MS, accurate mass was verified and optimal signal was determined from the integrated area under the curve for the extracted ion chromatogram of each metabolite adduct, mobile phase pH, and polarity (increasing FIA-MS signal, white to magenta Heatmap). The optimal FIA-MS signal conditions of each metabolite were manually filtered and binned to program an automated liquid handling method to construct the MIDAS metabolite pools (P1, P2, P3, and P4) according to the specific conditions of metabolite analysis by FIA-MS. **(C)** SDS-PAGE analysis of the purified proteins analyzed by MIDAS. mTORC1 regulators and the enzymes from central carbon metabolism are labeled.

**Figure S2.**
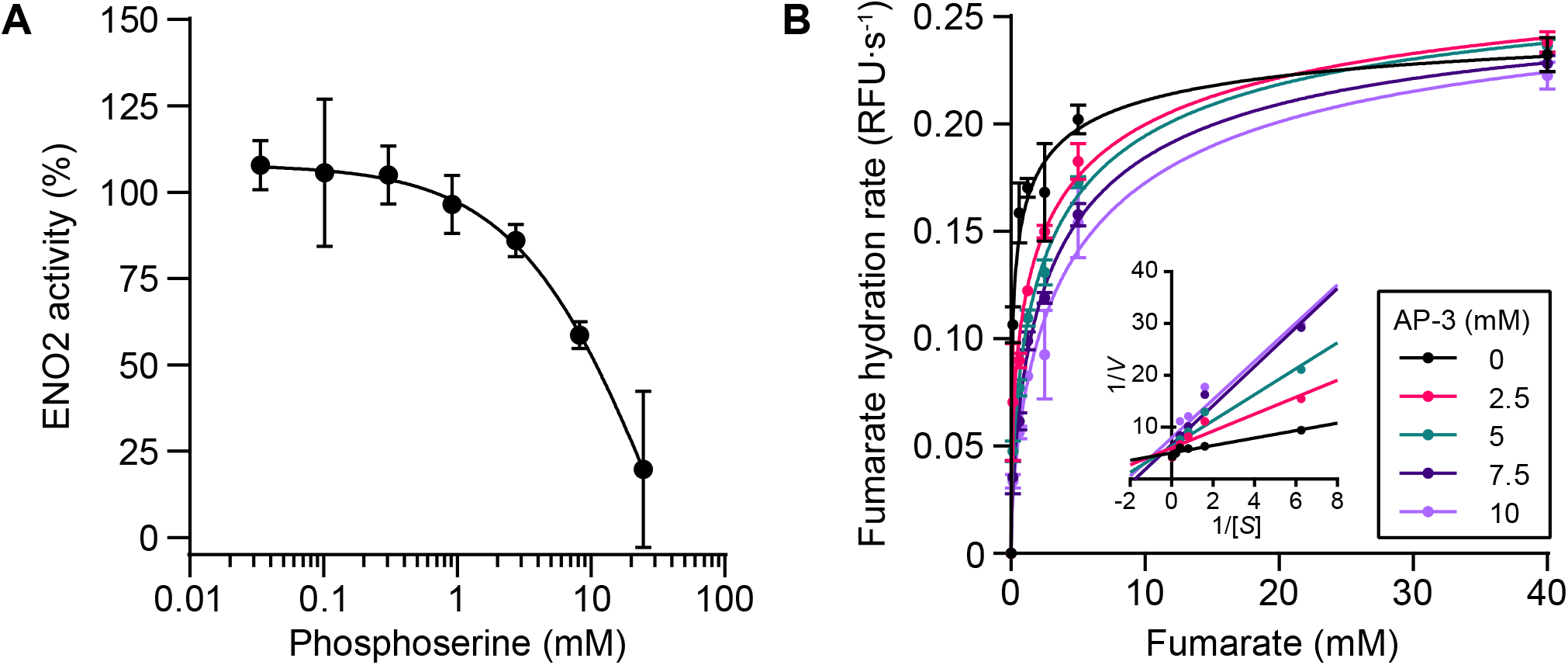
Enzymatic activities of enolase and fumarase with interacting metabolites. Activity of enolase, expressed as a percentage of vehicle control, was determined in the presence of varying concentrations of phosphoserine (pSer) using a coupled enzyme kinetic assay with pyruvate kinase/lactate dehydrogenase. The experiment was performed in triplicate and the mean ± SD are plotted. Relative fumarate hydration rate by fumarase was determined in the presence of varying concentrations of substrate, fumarate, and the inhibitor, 2-amino-3-phosphonopropionic acid (AP-3, colored) using a malate dehydrogenase coupled enzyme assay. (Inset) Lineweaver–Burk plot demonstrating competitive inhibition. The experiment was performed in triplicate and the mean ± SD are plotted.

**Figure S3.**
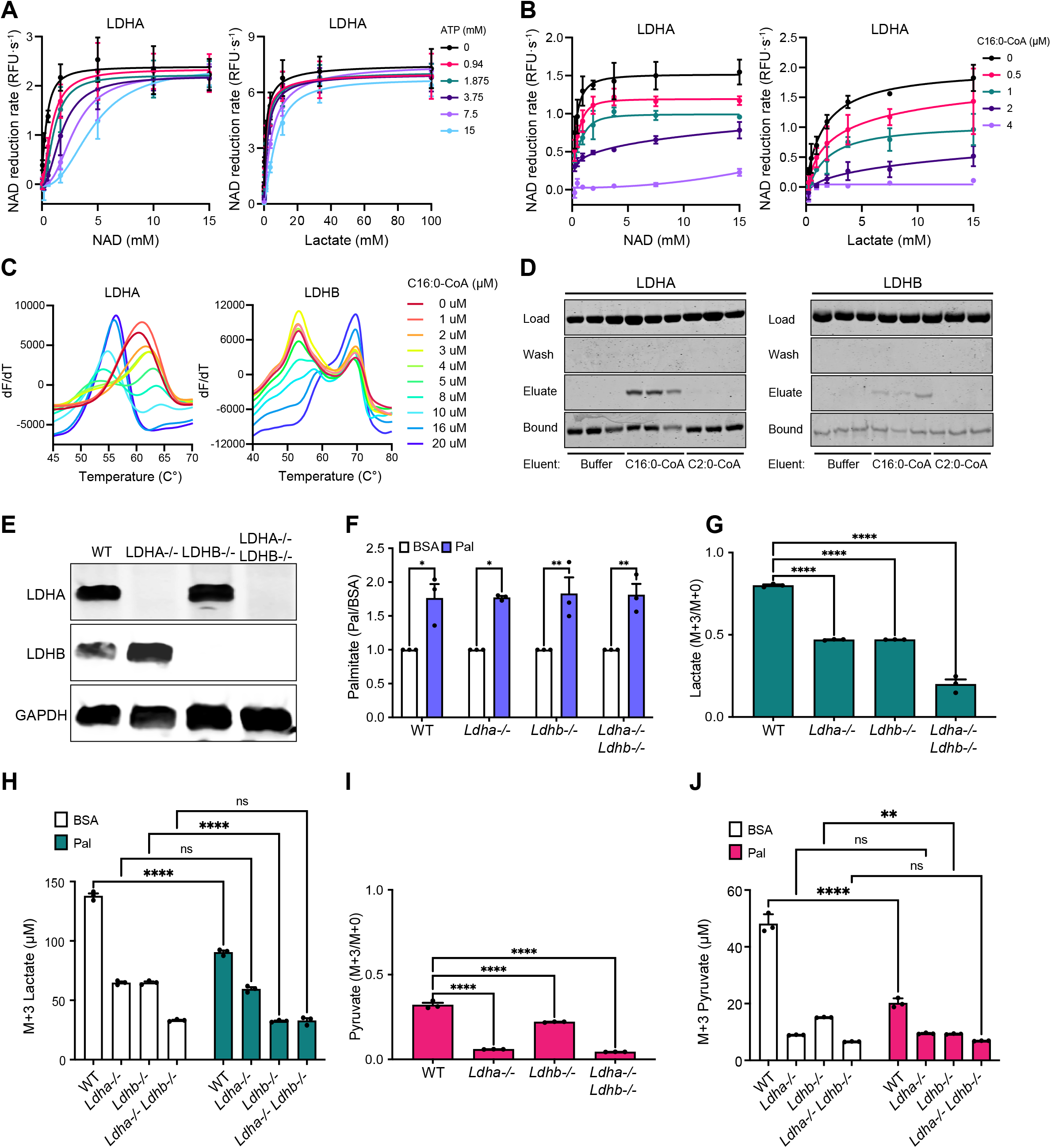
Lactate dehydrogenase interacts with and is inhibited by nucleotides and long-chain acyl-CoA. **(A)** Relative NAD reduction rate by LDHA was determined in the presence of varying concentrations of NAD (cofactor) or lactate (substrate), and ATP (colored) using a lactate dehydrogenase enzyme assay. The experiments were performed in triplicate and the mean ± SD are plotted. **(B)** Relative NAD reduction rate by LDHA was determined in the presence of varying concentrations of NAD (cofactor) or lactate (substrate), and palmitoyl-CoA (C16:0-CoA) (colored) using a lactate dehydrogenase enzyme assay. The experiments were performed in triplicate and the mean ± SD are plotted. **(C)** LDHA and LDHB were analyzed by PROTEOSTAT DSF in the presence of increasing concentrations of C16:0-CoA (colored). dF/dT was determined as a function of temperature. Representative experiments from triplicate experiments are presented. **(D)** Palmitoyl-CoA-Agarose pull-down assay with LDHA or LDHB treated with buffer control, palmitoyl-CoA (C16:0-CoA), or acetyl-CoA (C2:0-CoA) (Eluent). Protein input (Load), post-5x wash (Wash), concentrated supernatant post-eluent treatment (Eluate), protein bound to palmitoyl-CoA-agarose beads post-eluent treatment (Bound). The experiment was performed in triplicate. **(E)** Representative immunoblot of LDHA and LDHB in the indicated H9c2-derived cell lines. **(F)** Fold change of intracellular palmitate in *Ldha−/−*, *Ldhb−/−*, or *Ldha−/−;Ldhb−/−* H9c2 cell lines in response to treatment with palmitate-conjugated BSA (Pal) relative to BSA vehicle control (BSA). **(G)** Changes in ^13^C enrichment of extracellular lactate in *Ldha−/−*, *Ldhb−/−*, or *Ldha−/−;Ldhb−/−* H9c2 cell lines in response to treatment with palmitate-conjugated BSA. **(H)** Concentration of ^13^C-labelled extracellular lactate in *Ldha−/−*, *Ldhb−/−*, or *Ldha−/−;Ldhb−/−* H9c2 cell lines in response to BSA vehicle control (BSA) or palmitate-conjugated BSA (Pal) treatment. **(I)** Changes in intracellular ^13^C enrichment of pyruvate in *Ldha−/−*, *Ldhb−/−*, or *Ldha−/−;Ldhb−/−* H9c2 cell lines in response to treatment with palmitate-conjugated BSA. **(J)** Concentration of intracellular ^13^C-labelled pyruvate in *Ldha−/−*, *Ldhb−/−*, or *Ldha−/−;Ldhb−/−* H9c2 cell lines in response to BSA vehicle control (BSA) or palmitate-conjugated BSA (Pal) treatment. **(F – G)** All experiments were performed in triplicate. Data are presented as mean ± SD. *p < 0.05, **p < 0.01, ***p < 0.001, ****p < 0.0001, determined by one-way ANOVA and Sidak’s multiple comparison test.

**Figure S4.**
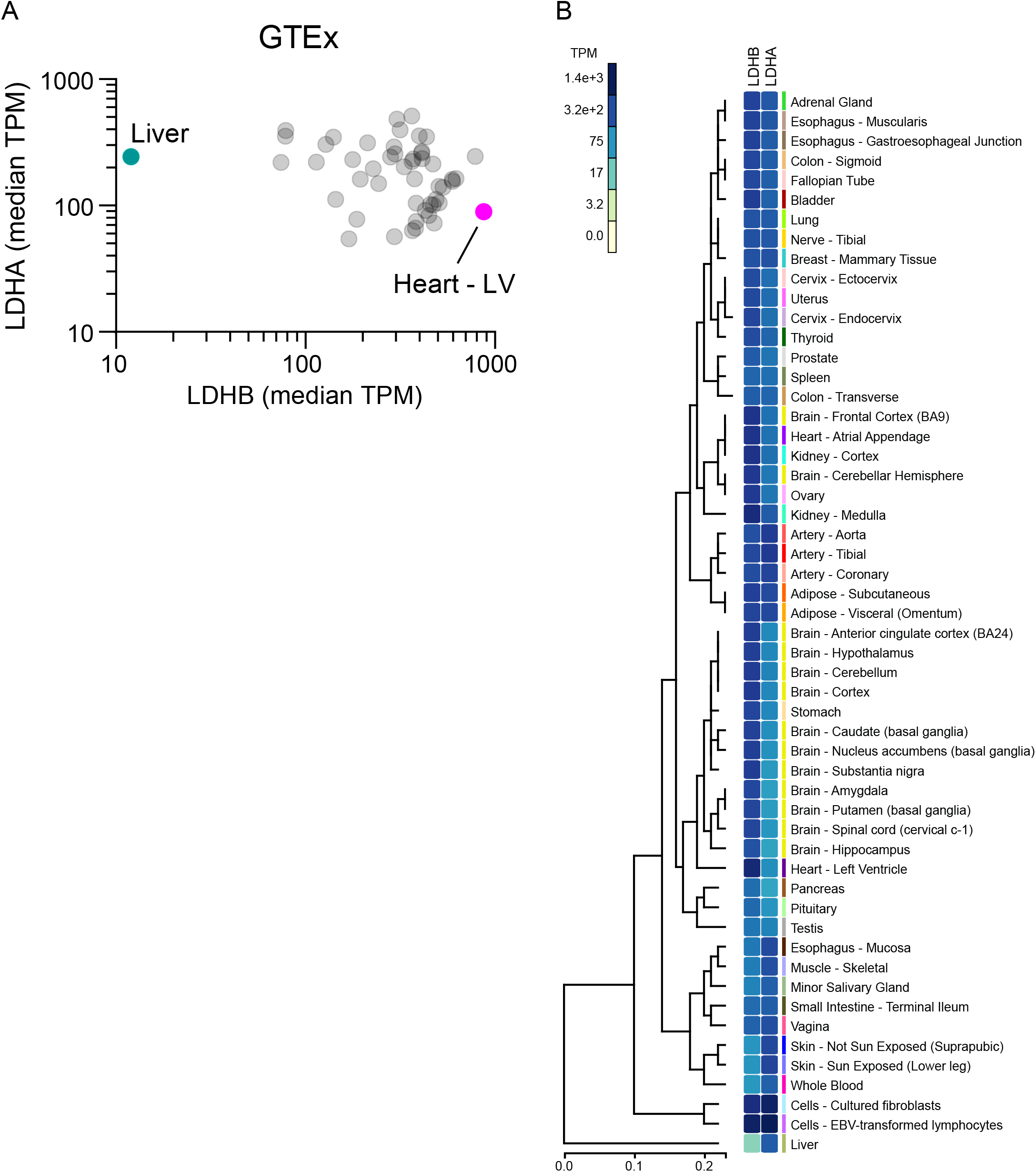
Tissues expression of lactate dehydrogenase segeragate in an isoform-specific manner. Differential gene expression of LDHA and LDHB across human tissues. (A) Scatter plot depicting median transcripts per million (TPM) for LDHA and LDHB. Note logarithmic axes. (B) Heatmap depicting median TPM for LDHA and LDHB across human tissues. Data obtained through GTEx Portal.

**Table S1.**
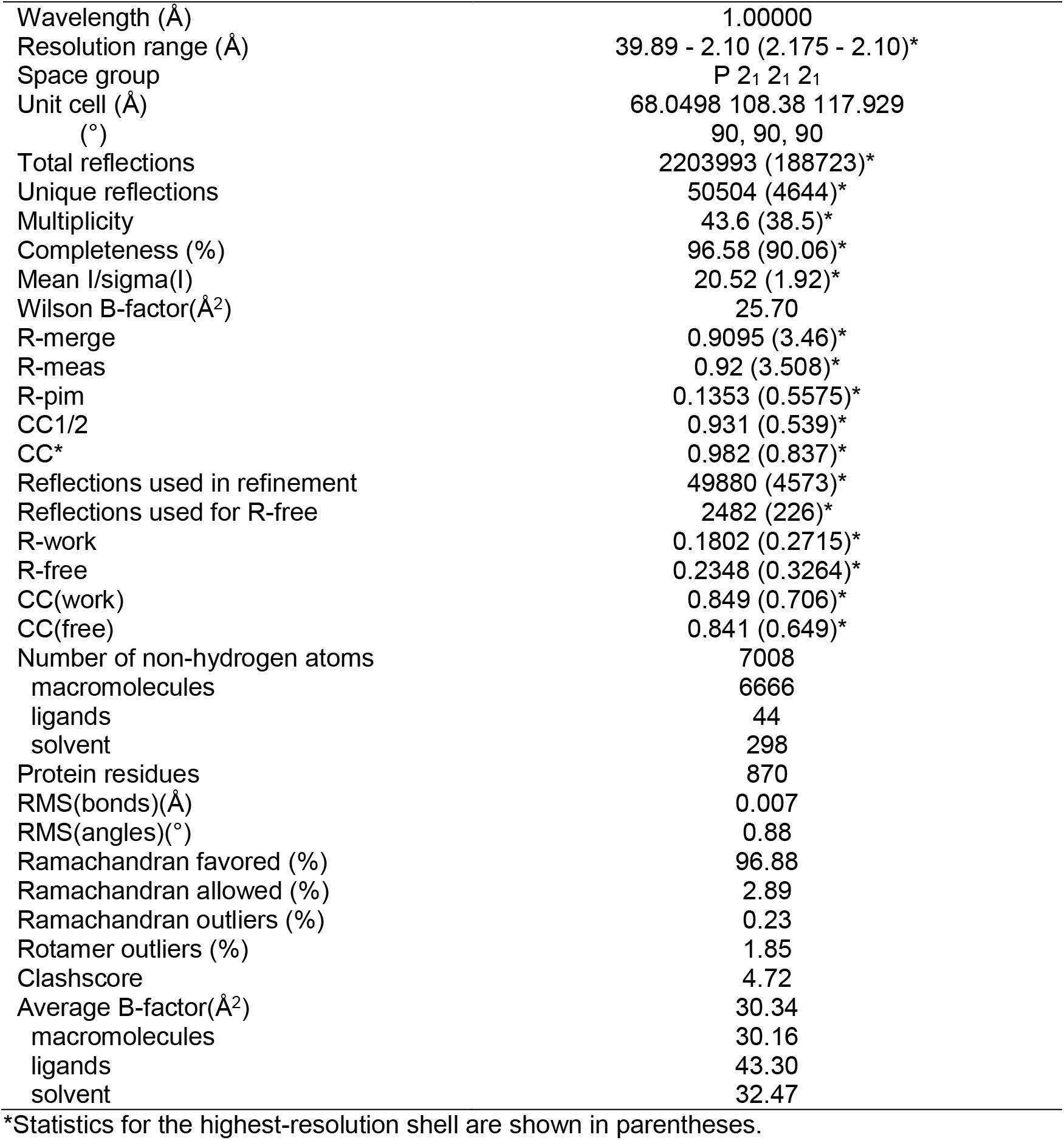
Phosphoserine-ENO2 data collection and refinement statistics.

|  |  |
| --- | --- |
| Wavelength (Å) | 1.00000 |
| Resolution range (Å) | 39.89 - 2.10 (2.175 - 2.10)* |
| Space group | P 2<sub>1</sub> 2<sub>1</sub> 2<sub>1</sub> |
| Unit cell (Å) | 68.0498 108.38 117.929 |
| (°) | 90, 90, 90 |
| Total reflections | 2203993 (188723)* |
| Unique reflections | 50504 (4644)* |
| Multiplicity | 43.6 (38.5)* |
| Completeness (%) | 96.58 (90.06)* |
| Mean I/sigma(I) | 20.52 (1.92)* |
| Wilson B-factor(Å<sup>2</sup>) | 25.70 |
| R-merge | 0.9095 (3.46)* |
| R-meas | 0.92 (3.508)* |
| R-pim | 0.1353 (0.5575)* |
| CC1/2 | 0.931 (0.539)* |
| CC* | 0.982 (0.837)* |
| Reflections used in refinement | 49880 (4573)* |
| Reflections used for R-free | 2482 (226)* |
| R-work | 0.1802 (0.2715)* |
| R-free | 0.2348 (0.3264)* |
| CC(work) | 0.849 (0.706)* |
| CC(free) | 0.841 (0.649)* |
| Number of non-hydrogen atoms | 7008 |
| macromolecules | 6666 |
| ligands | 44 |
| solvent | 298 |
| Protein residues | 870 |
| RMS(bonds)(Å) | 0.007 |
| RMS(angles)(°) | 0.88 |
| Ramachandran favored (%) | 96.88 |
| Ramachandran allowed (%) | 2.89 |
| Ramachandran outliers (%) | 0.23 |
| Rotamer outliers (%) | 1.85 |
| Clashscore | 4.72 |
| Average B-factor(Å<sup>2</sup>) | 30.34 |
| macromolecules | 30.16 |
| ligands | 43.30 |
| solvent | 32.47 |

\*Statistics for the highest-resolution shell are shown in parentheses.

**Table S2.** AP-3-fumarase data collection and refinement statistics.

|  |  |
| --- | --- |
| Wavelength (Å) | 1.323630 |
| Resolution range (Å) | 49.80 - 2.15 (2.20 - 2.15)* |
| Space group | P 6<sub>5</sub> 2 2 |
| Unit cell (Å) | 190.90, 116.24, 190.90 |
| (°) | 90, 120, 90 |
| Total reflections | 2660677 (178767)* |
| Unique reflections | 67921 (4478)* |
| Multiplicity | 39.2 (39.9)* |
| Completeness (%) | 100 (100)* |
| Mean I/sigma(I) | 15.0 (1.1)* |
| Wilson B-factor | 43.3 |
| R-merge | 0.25 (4.82)* |
| R-meas | 0.26 (4.95)* |
| R-pim | 0.056 (1.088)* |
| CC1/2 | 0.999 (0.458)* |
| CC* | - |
| Reflections used in refinement | 67877 |
| Reflections used for R-free | 3399 |
| R-work | 0.1858(0.3100)* |
| R-free | 0.2088(0.3300)* |
| CC(work) | 0.965(0.6406)* |
| CC(free) | 0.955(0.6746)* |
| Number of non-hydrogen atoms | 7191 |
| macromolecules | 6847 |
| ligands | 44 |
| solvent | 300 |
| Protein residues | 922 |
| RMS deviations (bonds) | 0.0031 |
| RMS deviations (angles) | 1.266 |
| Ramachandran favored (%) | 96.82 |
| Ramachandran allowed (%) | 3.18 |
| Ramachandran outliers (%) | 0.00 |
| Rotamer outliers (%) | 0.84 |
| Clashscore | 2.92 |
| Average B-factor | 52.00 |
| macromolecules | 52.09 |
| ligands | 56.03 |
| solvent | 50.97 |

\*Statistics for the highest-resolution shell are shown in parentheses.

**Other Supplementary Materials for this manuscript include the following:**

**Data S1. MIDAS metabolite library**

**Data S2. FIA-MS properties of MIDAS metabolites**

**Data S3. MIDAS proteins**

**Data S4. MIDAS protein-metabolite interactions**

## References

1. E. L. Huttlin et al., The BioPlex Network: A Systematic Exploration of the Human Interactome. Cell 162, 425–440 (2015).

2. D. S. Gilmour, J. T. Lis, Detecting protein-DNA interactions in vivo: distribution of RNA polymerase on specific bacterial genes. Proc Natl Acad Sci U S A 81, 4275–4279 (1984).

3. D. S. Gilmour, J. T. Lis, In vivo interactions of RNA polymerase II with genes of Drosophila melanogaster. Mol Cell Biol 5, 2009–2018 (1985).

4. M. Diether U. Sauer, Towards detecting regulatory protein-metabolite interactions. Curr Opin Microbiol 39, 16–23 (2017).

5. T. Orsak et al., Revealing the allosterome: systematic identification of metabolite-protein interactions. Biochemistry 51, 225–232 (2012).

6. L. Chantranupong et al., The CASTOR Proteins Are Arginine Sensors for the mTORC1 Pathway. Cell 165, 153–164 (2016).

7. R. L. Wolfson et al., Sestrin2 is a leucine sensor for the mTORC1 pathway. Science 351, 43–48 (2016).

8. K. Inoki, Y. Li, T. Xu, K. L. Guan, Rheb GTPase is a direct target of TSC2 GAP activity and regulates mTOR signaling. Genes Dev 17, 1829–1834 (2003).

9. P. Sun, Y. Liu, T. Ma, J. Ding, Structure and allosteric regulation of human NAD-dependent isocitrate dehydrogenase. Cell Discov 6, 94 (2020).

10. B. Chaneton et al., Serine is a natural ligand and allosteric activator of pyruvate kinase M2. Nature 491, 458–462 (2012).

11. R. Lopez-Alemany et al., Inhibition of cell surface mediated plasminogen activation by a monoclonal antibody against alpha-Enolase. Am J Hematol 72, 234–242 (2003).

12. S. Feo, D. Arcuri, E. Piddini, R. Passantino, A. Giallongo, ENO1 gene product binds to the c-myc promoter and acts as a transcriptional repressor: relationship with Myc promoter-binding protein 1 (MBP-1). FEBS Lett 473, 47–52 (2000).

13. T. Weaver, L. Banaszak, Crystallographic studies of the catalytic and a second site in fumarase C from Escherichia coli. Biochemistry 35, 13955–13965 (1996).

14. K. Nakahigashi et al., Systematic phenome analysis of Escherichia coli multiple-knockout mutants reveals hidden reactions in central carbon metabolism. Mol Syst Biol 5, 306 (2009).

15. S. Galvan-Pena et al., Malonylation of GAPDH is an inflammatory signal in macrophages. Nat Commun 10, 338 (2019).

16. M. Nakano, S. Funayama, M. B. de Oliveira, S. L. Bruel, E. M. Gomes, D-glyceraldehyde-3-phosphate dehydrogenase from HeLa cells--1. Purification and properties of the enzyme. Comp Biochem Physiol B 102, 873–877 (1992).

17. M. Oguchi, E. Gerth, B. Fitzgerald, J. H. Park, Regulation of glyceraldehyde 3-phosphate dehydrogenase by phosphocreatine and adenosine triphosphate. IV. Factors affecting in vivo control of enzymatic activity. J Biol Chem 248, 5571–5576 (1973).

18. M. Yuan et al., An allostatic mechanism for M2 pyruvate kinase as an amino-acid sensor. Biochem J 475, 1821–1837 (2018).

19. K. Ashizawa, P. McPhie, K. H. Lin, S. Y. Cheng, An in vitro novel mechanism of regulating the activity of pyruvate kinase M2 by thyroid hormone and fructose 1, 6-bisphosphate. Biochemistry 30, 7105–7111 (1991).

20. S. Hui et al., Glucose feeds the TCA cycle via circulating lactate. Nature 551, 115–118 (2017).

21. E. K. Wiese et al., Enzymatic activation of pyruvate kinase increases cytosolic oxaloacetate to inhibit the Warburg effect. Nat Metab 3, 954–968 (2021).

22. G. J. Gowans, D. G. Hardie, AMPK: a cellular energy sensor primarily regulated by AMP. Biochem Soc Trans 42, 71–75 (2014).

23. T. R. Koves et al., Mitochondrial overload and incomplete fatty acid oxidation contribute to skeletal muscle insulin resistance. Cell Metab 7, 45–56 (2008).

24. D. K. Sindelar et al., The role of fatty acids in mediating the effects of peripheral insulin on hepatic glucose production in the conscious dog. Diabetes 46, 187–196 (1997).

25. J. Billiard et al., Quinoline 3-sulfonamides inhibit lactate dehydrogenase A and reverse aerobic glycolysis in cancer cells. Cancer Metab 1, 19 (2013).

26. G. Rai et al., Discovery and Optimization of Potent, Cell-Active Pyrazole-Based Inhibitors of Lactate Dehydrogenase (LDH). J Med Chem 60, 9184–9204 (2017).

27. G. A. Bezerra et al., Crystal structure and interaction studies of human DHTKD1 provide insight into a mitochondrial megacomplex in lysine catabolism. IUCrJ 7, 693–706 (2020).

28. G. A. Bezerra et al., Identification of small molecule allosteric modulators of 5,10-methylenetetrahydrofolate reductase (MTHFR) by targeting its unique regulatory domain. Biochimie 183, 100–107 (2021).

29. Q. Hao et al., Sugar phosphate activation of the stress sensor eIF2B. Nat Commun 12, 3440 (2021).

30. J. D. Storey, R. Tibshirani, Statistical significance for genomewide studies. Proc Natl Acad Sci U S A 100, 9440–9445 (2003).

31. F. H. Niesen, H. Berglund, M. Vedadi, The use of differential scanning fluorimetry to detect ligand interactions that promote protein stability. Nat Protoc 2, 2212–2221 (2007).

32. N. Satani et al., ENOblock Does Not Inhibit the Activity of the Glycolytic Enzyme Enolase. PLoS One 11, e0168739 (2016).

33. T. G. Battye, L. Kontogiannis O. Johnson, H. R. Powell, A. G. Leslie, iMOSFLM: a new graphical interface for diffraction-image processing with MOSFLM. Acta Crystallogr D Biol Crystallogr 67, 271–281 (2011).

34. P. Evans, Scaling and assessment of data quality. Acta Crystallogr D Biol Crystallogr 62, 72–82 (2006).

35. P. R. Evans, G. N. Murshudov, How good are my data and what is the resolution? Acta Crystallogr D Biol Crystallogr 69, 1204–1214 (2013).

36. A. J. McCoy et al., Phaser crystallographic software. J Appl Crystallogr 40, 658–674 (2007).

37. D. Liebschner et al., Macromolecular structure determination using X-rays, neutrons and electrons: recent developments in Phenix. Acta Crystallogr D Struct Biol 75, 861–877 (2019).

38. P. Emsley, B. Lohkamp, W. G. Scott, K. Cowtan, Features and development of Coot. Acta Crystallogr D Biol Crystallogr 66, 486–501 (2010).

39. M. A. Ajalla Aleixo, V. L. Rangel, J. K. Rustiguel, R. A. P. de Padua, M. C. Nonato, Structural, biochemical and biophysical characterization of recombinant human fumarate hydratase. FEBS J 286, 1925–1940 (2019).

40. W. Kabsch, Xds. Acta Crystallogr D Biol Crystallogr 66, 125–132 (2010).

41. A. Vagin A. Teplyakov, Molecular replacement with MOLREP. Acta Crystallogr D Biol Crystallogr 66, 22–25 (2010).

42. G. N. Murshudov, A. A. Vagin, E. J. Dodson, Refinement of macromolecular structures by the maximum-likelihood method. Acta Crystallogr D Biol Crystallogr 53, 240–255 (1997).

43. C. J. Williams et al., MolProbity: More and better reference data for improved all-atom structure validation. Protein Sci 27, 293–315 (2018).

44. A. Xiao et al., CasOT: a genome-wide Cas9/gRNA off-target searching tool. Bioinformatics 30, 1180–1182 (2014).

45. A. A. Cluntun et al., The rate of glycolysis quantitatively mediates specific histone acetylation sites. Cancer Metab 3, 10 (2015).

46. M. J. Lukey et al., Liver-Type Glutaminase GLS2 Is a Druggable Metabolic Node in Luminal-Subtype Breast Cancer. Cell Rep 29, 76–88 e77 (2019).

47. A. A. Cluntun et al., The pyruvate-lactate axis modulates cardiac hypertrophy and heart failure. Cell Metab 33, 629–648 e610 (2021).

48. J. M. Buescher et al., A roadmap for interpreting (13)C metabolite labeling patterns from cells. Curr Opin Biotechnol 34, 189–201 (2015).

49. J. Qin, G. Chai, J. M. Brewer, L. L. Lovelace, L. Lebioda, Structures of asymmetric complexes of human neuron specific enolase with resolved substrate and product and an analogous complex with two inhibitors indicate subunit interaction and inhibitor cooperativity. J Inorg Biochem 111, 187–194 (2012).

